# p24-Tango1 interactions ensure ER-Golgi interface stability and efficient transport

**DOI:** 10.1101/2024.02.23.580604

**Authors:** Ke Yang, Zhi Feng, José C. Pastor-Pareja

## Abstract

The eukaryotic p24 family, consisting of α-, β-, γ- and δ-p24 subfamilies, has long been known to be involved in regulating secretion. Despite increasing interest in these proteins, fundamental questions remain about their role. Here, we systematically investigated *Drosophila* p24 proteins. We discovered that members of all four p24 subfamilies are required for general secretion, and that their localizations between ER exit site (ERES) and Golgi are interdependent in an α→βδ→γ sequence. We also found that localization of p24 proteins and ERES determinant Tango1 requires interaction through their respective GOLD and SH3 lumenal domains, with Tango1 loss sending p24 proteins to the plasma membrane and vice versa. Finally, we show that p24 loss expands the COPII zone at ERES and increases the number of ER-Golgi vesicles, supporting a restrictive role of p24 proteins on vesicle budding for efficient transport. Our results reveal Tango1-p24 interplay as central to the generation of a stable ER-Golgi interface.

**Summary:** Yang et al. systematically analyze in *Drosophila* the function of the four p24 protein subfamilies and discover that interaction with Tango1 is essential for their concentration between ER and Golgi and for efficiency of COPII-mediated general secretory transport.

## INTRODUCTION

Efficient trafficking of secretory cargos from the endoplasmic reticulum (ER) to the Golgi apparatus is essential for the physiological health and the correct organization of eukaryotic cells. In the eukaryotic secretory pathway, cargos are collected at specialized regions of the ER called ER exit sites (ERES), from where they are transported to the Golgi with the assistance of the COPII vesicle budding machinery (Bannykh et al., 1996; Barlowe and Miller, 2013; Brandizzi and Barlowe, 2013; Zanetti et al., 2012). In addition to this forward secretory traffic, ERES concentrate as well the income of proteins and membranes traveling in the opposite direction from the Golgi to the ER (Lerich et al., 2012; Roy Chowdhury et al., 2020; Yang et al., 2021). ERES, therefore, are critical traffic junctions mediating both anterograde and retrograde transport. To do this, cells must bring together in the reduced space between ERES and Golgi the numerous cytoplasmic components of the different transport machineries and their multiple regulators. The question of how cells organize and maintain a dynamic but stable ER-Golgi interface for efficient transport in the face of constant forward and reverse membrane traffic has sparked great interest among cell biologists.

p24 proteins are a family of type-I transmembrane proteins highly conserved among eukaryotes. Identified as major constituents of both COPI and COPII vesicles (Otte et al., 2001; Schimmoller et al., 1995; Sohn et al., 1996; Stamnes et al., 1995), they are long known to be involved in secretion (Kaiser, 2000; Pastor-Cantizano et al., 2016). Based on sequence homology, p24 proteins are classified into four subfamilies: α-, β-, γ- and δ-p24 (Dominguez et al., 1998; Pastor-Cantizano et al., 2016; Strating et al., 2009). p24 proteins of all four subfamilies display a common modular structure (Fig. 1 A) consisting of a cleavable signal peptide, a luminal part with a Golgi dynamics (GOLD) domain (Anantharaman and Aravind, 2002), a single hydrophobic transmembrane region and a short cytosolic tail that contains well-characterized COPI and COPII recruiting motifs responsible for their cycling between the ER and Golgi (Dominguez et al., 1998; Fiedler et al., 1996). Given the striking conservation of their four subfamilies, abundant presence at the ER-Golgi interface and multiple disease connections (Roberts and Satpute-Krishnan, 2023), understanding the role of p24 proteins has been a prominent research goal in the secretion field for over two decades. However, despite a large number of studies and spiking interest in recent years, fundamental questions about them remain largely unresolved.

**Figure 1.**
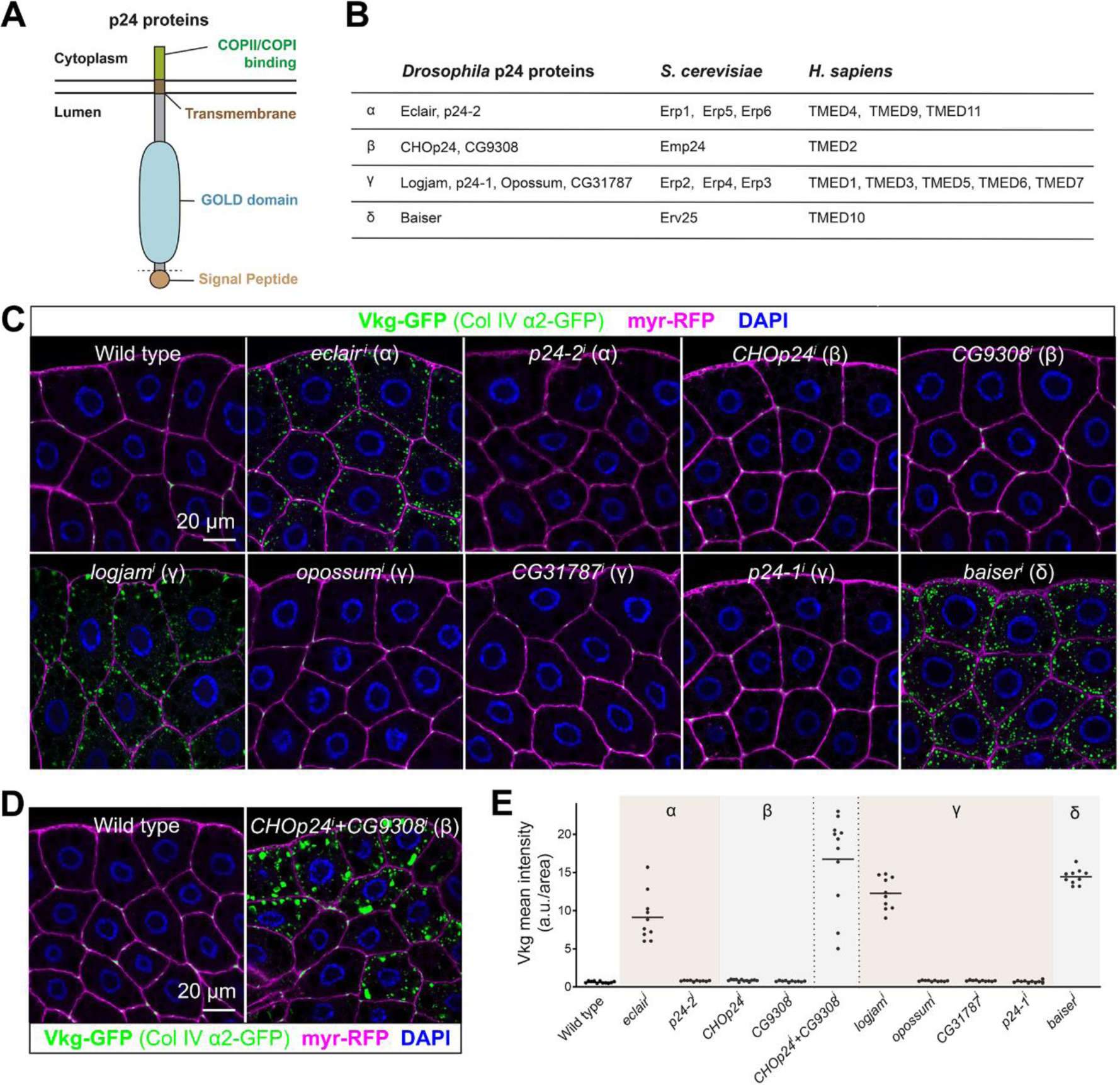
α-, β-, γ- and δ-p24 proteins are required for Collagen IV secretion in *Drosophila*. **(A)** Schematic domain organization of p24 family proteins, representing their cleavable secretion signal peptide, lumenal GOLD domain, transmembrane region and short cytoplasmic tail containing COPII/COPI recruitment motifs. **(B)** Classification into α, β, γ and δ subfamilies of p24 proteins in the fruit fly (*Drosophila melanogaster*), baker’s yeast (*Saccharomyces cerevisiae*) and humans (*Homo sapiens*). **(C, D)** Confocal images of L3 fat body adipocytes from wild type larvae and larvae where p24-encoding genes have been knocked down under control of fat body driver *BM-40-SPARC-GAL4* individually (C, *BM- 40-SPARC>eclair^i^*, *>p24-1^i^*, *>CHOp24^i^*, *>CG9308^i^*, *>logjam^i^*, *>opossum^i^*, *>CG31787^i^*, *>p24-1^i^* and *>baiser^i^*) and, for β-p24 proteins, in combination (D, *BM-40-SPARC>CHOp24^i^+CG9308^i^*), showing localization of Collagen IV (α2 chain Vkg-GFP in green). Plasma membrane labeled with GAL4-driven myr-RFP (magenta). Nuclei stained with DAPI (blue). **(E)** Quantification of intracellular Collagen IV retention measured from images like those in C and D. Each dot represents a measurement in one cell (n ≥ 10 per group). Horizontal lines indicate mean values. See also Figure S1.

p24 proteins are widely believed to function as specific cargo-interacting receptors for a collection of different protein cargos. These include GPI-anchored proteins (Bernat-Silvestre et al., 2020; Bonnon et al., 2010; Castillon et al., 2011; Manzano-Lopez et al., 2015; Muñiz et al., 2000; Takida et al., 2008), Wnt family ligands (Buechling et al., 2011; Port et al., 2011), G-protein coupled receptors (Luo et al., 2007), fibronectins (Hou and Jerome-Majewska, 2018), plant myrosinase-associated protein GLL23 (Jancowski et al., 2014), insulin (Hosaka et al., 2007; Zhang and Volchuk, 2010), Rac-GAP chimaerin (Wang and Kazanietz, 2002), Toll-like receptor 4 (Liaunardy-Jopeace et al., 2014), and, more recently, leaderless cargos such as Interleukin-1 (Zhang et al., 2020). Furthermore, defective function of p24 proteins is associated with Alzheimer’s disease as mediators of APP trafficking modulating γ-secretase cleavage (Chen et al., 2006; Hasegawa et al., 2010; Vetrivel et al., 2007). Nevertheless, along with reports of defects in the transport of specific cargos, evidence of more general impairments upon p24 deficiencies exist in the literature as well, such as altered Golgi-ER retrograde transport (Aguilera-Romero et al., 2008; Gommel et al., 2001; Majoul et al., 1998; Montesinos et al., 2014) and abnormal Golgi morphology (D’Arcangelo et al., 2015; Denzel et al., 2000; Koegler et al., 2010; Lavoie et al., 1999; Mitrovic et al., 2008; Pastor-Cantizano et al., 2018; Rojo et al., 2000). Moreover, broad roles have been ascribed to p24 proteins in mediating ER retention for quality control (Belden and Barlowe, 2001; Dvela-Levitt et al., 2019; Gomez-Navarro et al., 2020; Lopez et al., 2020; Ma et al., 2017; Springer et al., 2000; Wen and Greenwald, 1999), membrane contact during autophagosome formation from the ERGIC (ER Golgi Intermediate Compartment) (Li et al., 2022) and lipid transfer between the ER and Golgi (Anwar et al., 2022). Fitting all these proposed functions, cargo-specific and general, into a consistent view is problematic. Furthermore, p24 proteins have been reported to function in heteromeric complexes (Fullekrug et al., 1999; Marzioch et al., 1999) and the wide conservation of the four subfamilies hints at important, non-redundant roles in the secretory pathway; however, because of their similar organization and structure, even when members of different subfamilies are compared, the question of whether they play differentiated or redundant roles remains unanswered. Complicating analysis of these issues through genetics, yeast p24 mutants are viable and show only mild defects, even when combined into an octuple mutant where all members of the four p24 subfamilies are deleted (Springer et al., 2000).

The genome of the fruit fly *Drosophila melanogaster* encodes nine proteins of the p24 family, distributed among the four conserved subfamilies as follows (Fig. 1 B): Eclair and p24-2 belong to the α-p24 subfamily; CHOp24 and CG9308 to the β-p24 subfamily; Logjam, CG31787, Opossum and p24-1 are γ-24 subfamily members, and, finally, Baiser is the only δ-p24 subfamily representative (Carney and Bowen, 2004). In contrast to the situation in yeast, *Drosophila* p24 proteins play clearly essential roles, ubiquitous knock down of each of them in all tissues producing lethality or severely reduced viability (Saleem et al., 2012). Phenotypic loss of function analysis of *Drosophila* p24 proteins has revealed defects in embryonic patterning (Bartoszewski et al., 2004), oviposition (Boltz et al., 2007; Carney and Taylor, 2003), fecundity and male fertility (Saleem et al., 2012), and the stress response (Boltz and Carney, 2008). In addition, detailed mechanistic studies concluded that p24 proteins interact with Wingless and other WNT family ligands, and are required for their secretion (Buechling et al., 2011; Port et al., 2011; Zang et al., 2015), raising again the question of whether p24 proteins act in the early secretory pathway as specific receptors for the transport of particular cargos. Furthermore, regarding the redundancy and relations among the different subfamilies, a systematic analysis of protein localization and mutual functional requirements has not been carried out.

*Drosophila* is a powerful model for investigating protein secretion and the early secretory pathway. Genetic screens in flies have identified conserved new secretory genes (Bard et al., 2006; Ke et al., 2018; Kondylis et al., 2011; Tiwari et al., 2015; Wendler et al., 2010). Additional advantages of researching secretion in *Drosophila* is the availability of sophisticated tools for transgenic tagging and tissue-specific functional interrogation. Many recent studies have taken advantage of these to dissect secretory traffic in an animal (Fujii et al., 2020; Glashauser et al., 2023; Johnson et al., 2020; Ma et al., 2020; Park et al., 2022; Song et al., 2022; van Leeuwen et al., 2018; Zajac and Horne-Badovinac, 2022; Zhou et al., 2023). In *Drosophila*, the early secretory pathway is organized into secretory units, tens to hundreds per cell, in which ERES lie in close proximity to Golgi ministacks (Kondylis and Rabouille, 2009). We have previously characterized the organization of these ERES-Golgi units using 3D-SIM (Structured Illumination Microsocopy), TEM (Transmission Electron Microscopy) and FIB-SEM (Focused Ion Beam-Scanning Electron Microscopy) (Yang et al., 2021). Besides occasional continuities between ERES and pre-*cis*-Golgi, we could distinguish two populations of vesicles at the ER-Golgi interface: one at the center of the ERES cup, corresponding to the highest COPII concentration, and the other in the periphery, consistent in size and localization with retrograde COPI vesicles (Yang et al., 2021). A critical protein in the maintenance of this ER-Golgi interface is Tango1 (Transport and Golgi organization 1), an ERES-localized transmembrane protein discovered in a screening in *Drosophila* S2 cells (Bard et al., 2006; Saito et al., 2009). Tango1 is the single *Drosophila* member of the MIA/cTAGE family, only present in animals (Feng et al., 2020). Loss of Tango1 function has been shown to impair secretion of multiple cargos in all examined *Drosophila* tissues (Lerner et al., 2013; Liu et al., 2017; Pastor-Pareja and Xu, 2011; Reynolds et al., 2019; Rios-Barrera et al., 2017; Zhang et al., 2014). In absence of Tango1, ERES become smaller and detach from Golgi (Liu et al., 2017), indicating that Tango1, among other roles, can function as a tether and organizer of the ER-Golgi interface (Feng et al., 2020; McCaughey et al., 2021; Saito and Maeda, 2019). The cytoplasmic part of Tango1, capable of self-interacting (Liu et al., 2017), may have a chief role in this organizing function, whereas the role of the ER lumenal part of Tango1, which contains an SH3 domain reported to bind cargos directly or through adaptors (Arnolds and Stoll, 2023; Ishikawa et al., 2016; Saito et al., 2009; Yuan et al., 2018), is less understood. Mechanisms that ensure concentration of Tango1 at ERES could be of prime importance to regulate their size and protect the stability of the ERES-Golgi interface.

Here, using the larval fat body as a screening system, we have carried out a systematic analysis of the p24 family in *Drosophila.* We show that presence of members of all four p24 subfamilies is necessary for general secretion and dissect their mutual requirements for localization between ERES and pre-cis-Golgi. We also show that p24 proteins and Tango1 interact in the ER lumen and mutually depend on each other for their localization at the ER-Golgi interface. Finally, our high-resolution analysis through FIB-SEM shows an excess of vesicles in p24 loss conditions. Overall, our results evidence that p24 proteins confer stability to the ER-Golgi interface by limiting COPII budding and preventing Tango1 scape from ERES.

## RESULTS

### All four p24 subfamilies are required for general secretion in *Drosophila*

In a previous screening, we found that *logjam*, encoding a *Drosophila* γ-p24 protein, is required for Collagen IV secretion by fat body adipocytes of the third instar larva (L3 stage) (Ke et al., 2018), the main source of Collagen IV for the basement membranes of the *Drosophila* larva (Pastor-Pareja and Xu, 2011). To better understand the role of Logjam and p24 proteins in the secretory pathway, we knocked down the expression of the remaining *Drosophila* p24 proteins in the fat body under control of the GAL4-UAS expression system and found that, same as *logjam* (*logjam^i^*), knock down of *eclair* (*eclair^i^*) and *baiser* (*baiser^i^*), respectively encoding α- and δ-p24 subfamily members, led to intracellular retention of Viking-GFP (Vkg-GFP), a functional GFP-trap fusion of the Collagen IV α2 chain (Fig. 1, C and E). According to RNAseq data from others (Chintapalli et al., 2007; Krause et al., 2022), *eclair* (α), *logjam* (γ) and *baiser* (δ) are highly expressed in the fat body (Fig. S1). While no defect was observed upon single knock down of *CHOp24* or *CG9308*, encoding the two *Drosophila* β-p24 subfamily members, their simultaneous knock down (*CHOp24^i^*+*CG9308^i^*) led to Vkg-GFP intracellular retention (Fig. 1, D and E), proving their intra-subfamily redundancy in the fat body. These results, altogether, show that the functions of members of the α-, β-, γ- and δ-p24 subfamilies are required in fat body adipocytes for efficient Collagen IV secretion.

p24 proteins are proposed to function as specific cargo receptors for certain kinds of proteins such as GPI-anchored proteins (Bonnon et al., 2010; Muñiz et al., 2000; Takida et al., 2008) or leaderless cargos (Zhang et al., 2020). Having shown their requirement in Collagen IV secretion, we decided to test their requirement in the transport of other cargos. We found that knock down in fat body adipocytes of *eclair* (α), *CHOp24+CG9308* (β), *logjam* (γ) or *baiser* (δ) caused defective secretion of not just Collagen IV (Fig. 2, A and B), but also of GPI-anchored GFP (GFP fused to GPI attachment signal from CD58) (Fig. 2, A and C), Apolipoprotein B-related Rfabg (Fig. 2, A and D), transmembrane protein CD8 (Fig. 2, A and E), and soluble secretion marker Secreted-GFP (GFP coupled to a signal peptide) (Fig. 2, A and F). Hence, similar to knock down of COPII coat component Sec31 (Fig. 2, A-F), knock down of p24 proteins caused defective secretion of all examined cargos. We additionally examined retrograde transport marker GFP-KDEL, which concentrates at fat body ERES as a result of Golgi-to-ER recycling by the retrograde KDEL receptor (KdelR) (Yang et al., 2021). In contrast with its clearance from the cell upon *KdelR* knock down (*KdelR^i^*), GFP-KDEL showed strong intracellular retention in the ER when we knocked down *eclair* (α), *CHOp24*+*CG9308* (β), *logjam* (γ) or *baiser* (δ) (Fig. 2 G), indicating a primary defect in ER-to-Golgi cargo trafficking. Based on these data, we conclude that p24 proteins of all four subfamilies are required in fat body adipocytes for efficient anterograde transport in the general secretory pathway.

**Figure 2.**
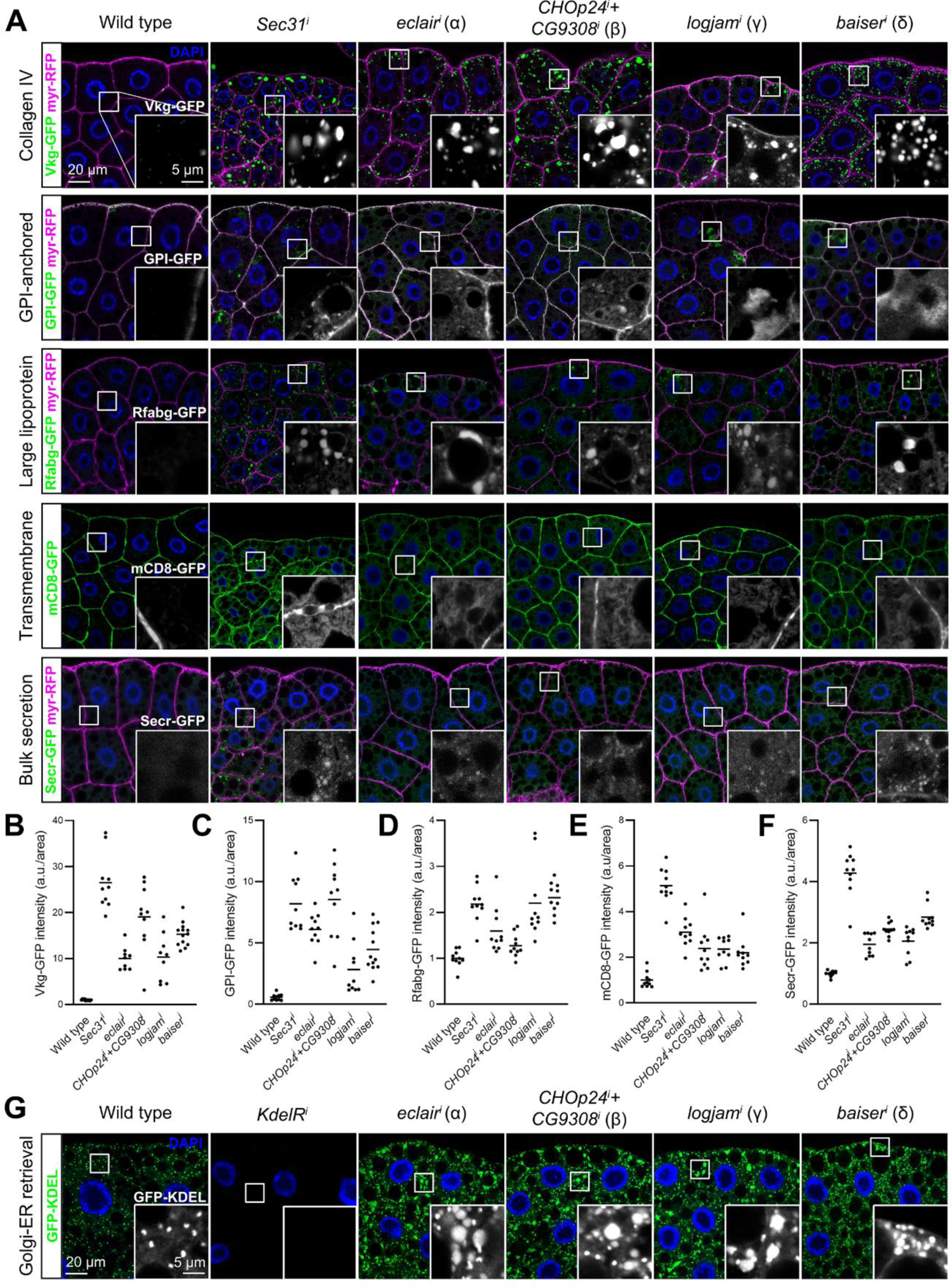
p24 proteins are required for general ER-Golgi transport. **(A)** Confocal images of L3 fat body adipocytes showing localization in green of Vkg-GFP (Collagen IV protein trap), GPI-GFP (driven by *Cg-GAL4*), Rfabg-GFP (driven by endogenous promoter), mCD8-GFP (driven by *BM-40-SPARC-GAL4*), and Secreted-GFP (driven by *BM-40-SPARC-GAL4*). Fat body was dissected from wild type larvae and larvae where genes encoding COPII coat component Sec31, α-p24 Eclair, β-p24 CHOp24 and CG9308, γ-p24 Logjam and δ-p24 Baiser had been knocked down under control of fat body drivers *Cg-GAL4* (for GPI-GFP) or *BM-40-SPARC-GAL4* (for Vkg-GFP, Rfabg-GFP, mCD8-GFP and Secr-GFP). Plasma membrane labeled with GAL4-driven myr-RFP (magenta), except for mCD8-GFP images. Nuclei stained with DAPI (blue). Magnified insets in the lower right corner of each image show isolated GFP signal in white. **(B-F)** Quantification of intracellular retention of Vkg-GFP (B), GPI-GFP (C), Rfabg-GFP (D), mCD8-GFP (E) and Secr-GFP (F), measured from images like those in A. Each dot represents a measurement in one cell (n > 10 per group). Horizontal lines indicate mean values. **(G)** Confocal images of L3 fat body adipocytes showing localization in green of GFP-KDEL (driven by *Cg-GAL4*). Fat body was dissected from wild type larvae and larvae where genes encoding KDEL receptor, Eclair, CHOp24+CG9308, Logjam and Baiser have been knocked down under control of *Cg-GAL4*. Nuclei stained with DAPI (blue). Magnified insets in the lower right corner of each image show isolated GFP signal in white.

### *Drosophila* p24 proteins concentrate between ERES and pre*-cis*-Golgi

To better understand the role of p24 proteins in the *Drosophila* secretory pathway, we investigated next their localization. In order to visualize p24 proteins, we added a GFP tag to the N-terminal of Eclair (α), CHOp24 (β), Logjam (γ) and Baiser (δ) after their signal peptides and expressed these tagged forms in fat body adipocytes. Through super resolution 3D-SIM, we observed that p24 proteins localized at ERES-Golgi units, concentrating between the ERES (marker Tango1) and Golgi (mid-Golgi marker Mannosidase II) (Fig. 3, A-D). To confirm this, we created transgenic flies in which we knocked-in an mCherry tag at the N-terminal of Logjam after its signal peptide using CRISPR/Cas9 technology and used this endogenous [mCherry]Logjam to study in detail its localization within ERES-Golgi units. To do this, we imaged Logjam and ERES marker Tango1 together with markers of different Golgi compartments (Yang et al., 2021): *trans*-Golgi marker GalT (Fig. 3, E-G), mid-Golgi marker ManII (Fig. 3, H-J), *cis*-Golgi marker GMAP (Fig. 3, K-M) and pre*-cis*-Golgi marker Grasp65 (Fig. 3, N-P; CRISPR/Cas9 Grasp65[GFP] knock-in). As evidenced by signal plot profiles and peak distance quantification, Logjam localization is distinct from those of GalT, ManII and GMAP, while its highest concentration is closer to pre-*cis*-Golgi Grasp65 (Fig. 3 l). Furthermore, of all examined markers, γ-p24 Logjam most closely resembled COPII coatomer Sec13 (Fig. 3, Q-T; CRISPR/Cas9 Sec13[GFP] knock-in), suggesting a close relation with the COPII vesicle budding machinery. Our data, therefore, place p24 protein localization at the ER-Golgi interface (Fig. 3 U), consistent with cycling between ERES and pre*-cis*-Golgi.

**Figure 3.**
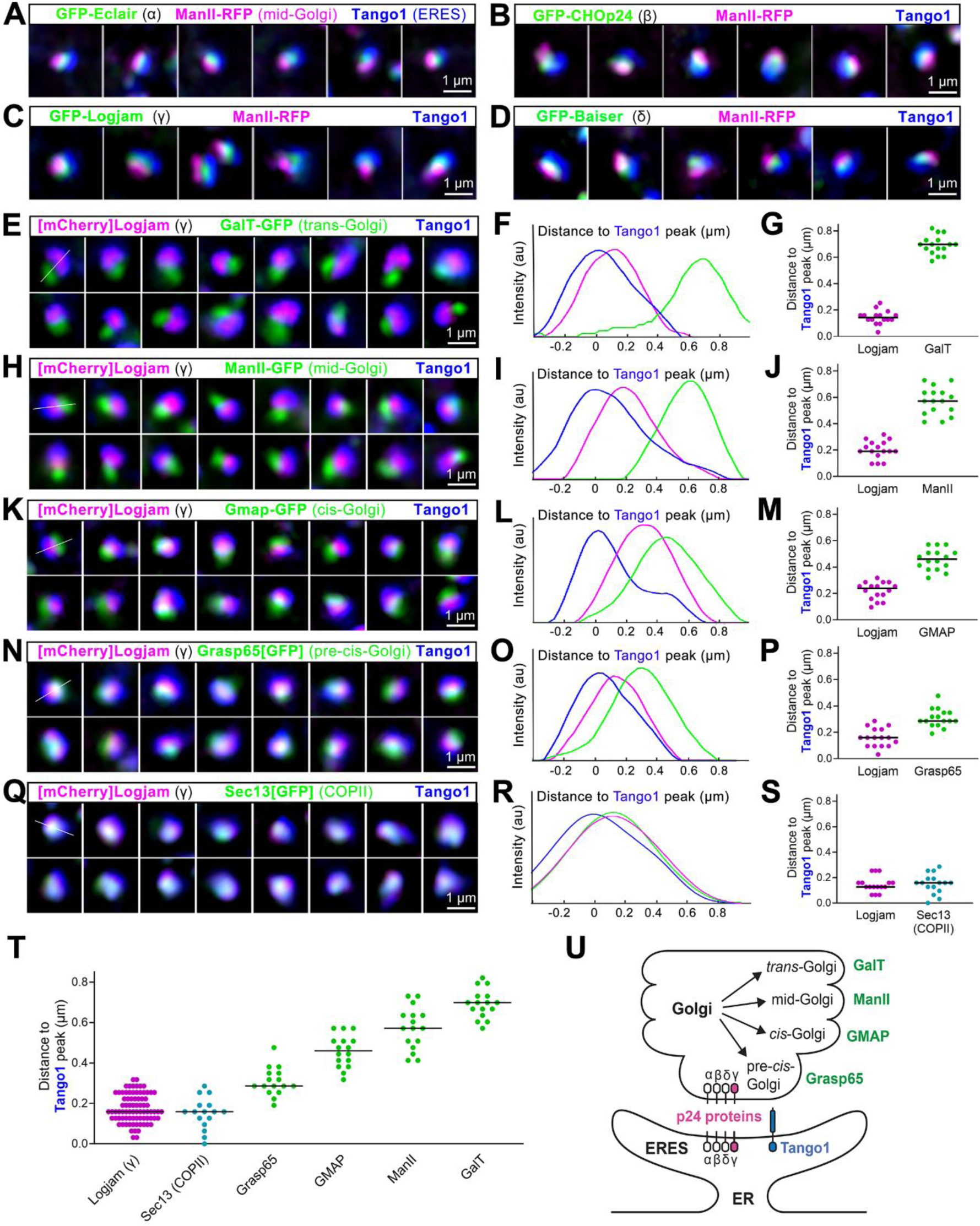
p24 proteins localize between ERES and pre-*cis*-Golgi. **(A-D)** Super resolution 3D-SIM images of ERES-Golgi units from L3 fat body adipocytes showing localization in green of GFP-tagged α-p24 Eclair (A), β-p24 CHOp24 (B), β-p24 Logjam (C) or δ-p24 Baiser (D), all driven by *Cg-GAL4*, mid-Golgi marker ManII (driven by *Cg-GAL4*, magenta) and ERES marker Tango1 (anti-Tango1, blue). **(E**, **H**, **K**, **N**, **Q)** Super resolution 3D-SIM images of ERES-Golgi units from L3 fat body adipocytes showing localization of endogenous γ-p24 Logjam ([mCherry]Logjam CRISPR/Cas9 knock-in, magenta) in relation to ERES Tango1 (anti-Tango1, blue) and, in green, *trans*-Golgi GalT-GFP (E, driven by *Cg-GAL4*), mid*-*Golgi ManII-GFP (H, driven by *Cg-GAL4*), *cis-*Golgi GMAP-GFP (K, protein trap), pre*-cis-*Golgi Grasp65[GFP] (N, CRISPR/Cas9 knock-in) and COPII coatomer Sec13[GFP] (Q, CRISPR/Cas9 knock-in). Images are maximum intensity projections of three to five sections (A-D, E, H, K, N and Q). **(F, I, L, O, R)** Signal profiles across individual ERES-Golgi units following the white lines in the upper left images in E, H, K, N and Q, respectively. **(G, J, M, P, S, T)** Graphs representing peak distances with respect to Tango1 in signal profiles like those in F, I, L, O and R, respectively. The horizontal lines indicate mean values. Each dot represents a measurement in one ERES-Golgi unit profile (G, J, M, P, S, n = 16 per group). Results summarized in T. **(U)** Schematic depiction of the localization of p24 proteins within an ERES-Golgi unit, as deduced from 3D-SIM image analysis.

### p24 protein localizations are interdependent in an α→βδ→γ sequence

p24 proteins have been reported to form heteromeric complexes (Fullekrug et al., 1999; Marzioch et al., 1999). Our finding that deficiency in each of the four p24 subfamilies resulted in defects in general secretion, and their similar localization at the ER-Golgi interface, led us to explore possible mutual requirements for their localization. To do this, we knocked down in the fat body the expression of *eclair* (α), *CHOp24*+*CG9308* (β), *logjam* (γ) or *baiser* (δ), and examined the effect of their loss in the localization of the remaining. In this way, we found that α-p24 Eclair concentration in ERES-Golgi units was unaffected by the loss of p24 proteins of the other subfamilies (Fig. 4, A and B). In contrast, localization of γ-p24 Logjam was defective when we knocked down the expression of members of each of the three other p24 subfamilies (Fig. 4, A and D), displaying a more diffuse ER distribution (Fig. 4 F). As for β-p24 CHOp24 and δ-p24 Baiser, their correct concentration depended on the presence not only of α-p24 Eclair, but also of each other (Fig. 4, A, C and E). Summarizing all these results together (Fig. 4 G), our analysis revealed an α→βδ→γ hierarchy for the correct localization of p24 proteins. In this hierarchy (Fig. 4 H), α-p24 is first to localize, independently, between ERES and pre-cis-Golgi, β- and δ-p24 are mutually dependent and dependent on the presence of α-p24, and, finally, γ-p24 is unable to concentrate in the absence of any of the other three.

**Figure 4.**
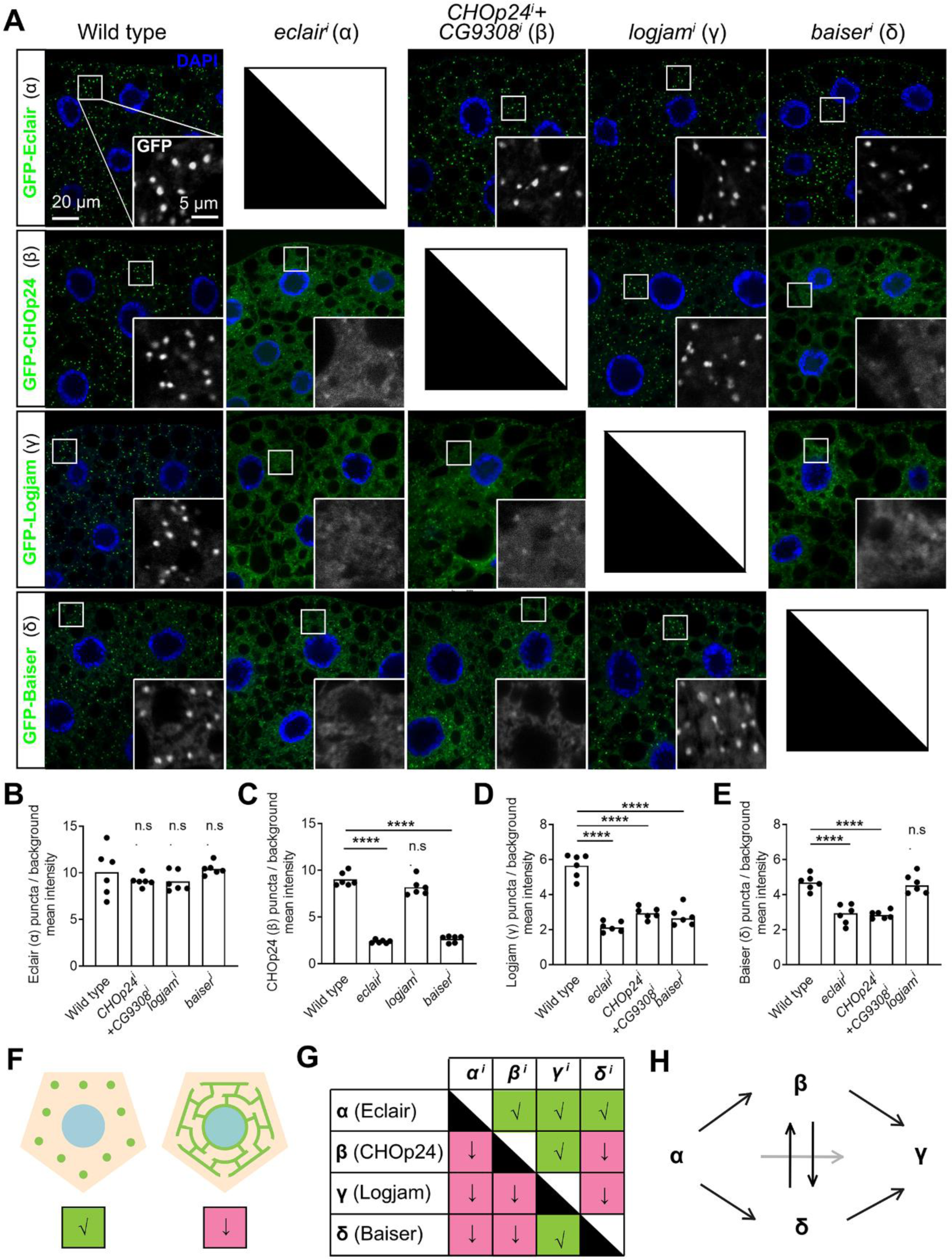
p24 protein localizations are interdependent in an α→βδ→γ sequence. **(A)** Confocal images of L3 fat body adipocytes showing localization in green of GFP-tagged α-p24 Eclair, β-p24 CHOp24, γ-p24 Logjam and δ-p24 Baiser, all driven by *Cg-GAL4*. Fat body was dissected from wild type larvae and larvae where genes encoding Eclair, CHOp24+CG9308, Logjam or Baiser had been knocked down under control of *Cg-GAL4.* Magnified insets in the lower right corner of each image show isolated GFP signal in white. Nuclei stained with DAPI (blue). **(B-E)** Graphs quantifying the effect on the localization of GFP-tagged Eclair (B), CHOp24 (C), Logjam (D) and Baiser (E) of the knock-down of indicated p24-encoding genes, measured from images like those in A. Graphs represent the ratio between the amounts of GFP signal concentrated in puncta and diffuse signal. Each dot represents a measurement from one cell (n = 6 in each group). Bar heights indicate mean value. *p* values from Brown-Forsythe ANOVA and Dunnett’s multiple comparisons tests (B, *p* = 0.7702 (n.s.) for *CHOp24^i^+CG9308^i^*, 0.7700 (n.s.) for *logjam^i^* and 0.9802 (n.s.) for *baiser^i^*), and one-way ANOVA and Dunnett’s multiple comparisons tests (C, *p* < 0.0001 (****) for *eclair^i^*, = 0.0735 (n.s.) for *logjam^i^*and < 0.0001 (****) for *baiser^i^*; D, *p* < 0.0001 (****) for *eclair^i^*, *CHOp24^i^+CG9308^i^* and *baiser^i^*; E, *p* < 0.0001 (****) for *eclair^i^* and *CHOp24^i^+CG9308^i^*, = 0.9846 (n.s.) for *logjam^i^*). **(F)** Illustration of punctate localization (√) and diffuse ER distribution (↓) observed for p24 proteins in A. **(G)** Summary of the effect of the knock-down of indicated p24-encoding genes on the localization of Eclair, CHOp24, Logjam and Baiser, according to B-E. **(H)** Model depicting requirements among p24 protein subfamilies for correct localization, as deduced from G.

### Localization of p24 proteins depends on a GOLD-SH3 interaction with Tango1

After observing dramatic changes in the localization of p24 proteins in our experiments, we proceeded to further investigate how p24 proteins maintain their steady localization at the ER-Golgi interface. To do that, we first knocked down in the fat body the expression of known *Drosophila* anterograde and retrograde ER-Golgi transport receptors encoded by *Ergic53* (*Ergic53^i^*) and *KdelR*, respectively, to explore whether they were involved in p24 localization, but found no difference in their normal punctate pattern when we imaged GFP-tagged versions of Eclair (α), CHOp24 (β), Logjam (γ) and Baiser (δ) (Fig. 5). Similarly, we observed no apparent defect in the localization of Eclair (α), CHOp24 (β), Logjam (γ) and Baiser (δ) upon knock down of *Grasp65* (*Grasp65^i^*; Fig. 5), encoding a protein of the pre-*cis*-Golgi required for secretion (Yang et al., 2021). In contrast to these, the distribution of p24 proteins of all four subfamilies strikingly changed upon knock down of ERES protein Tango1 (*Tango^i^*), showing diffuse ER localization and presence at the plasma membrane (Fig. 5). Endogenous γ-p24 Logjam tagged with mCherry displayed ER and plasma membrane mislocalization in *Tango1^i^* adipocytes as well (Fig. S2 A), confirming that Tango1 is required for the correct localization of p24 proteins.

**Figure 5.**
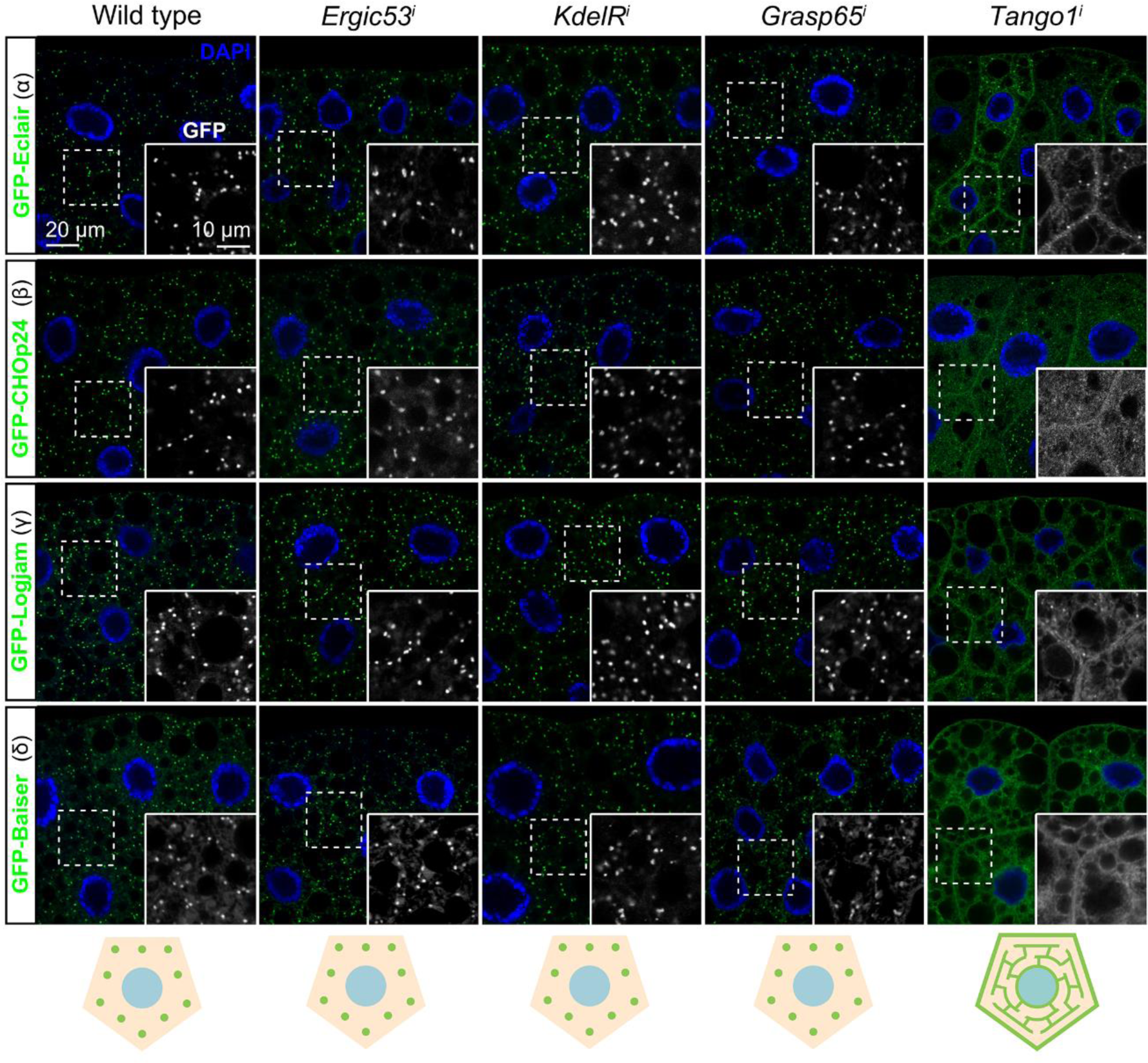
Concentration of p24 proteins at the ER-Golgi interface depends on Tango1. Confocal images of L3 fat body adipocytes showing localization in green of GFP-tagged α-p24 Eclair, β-p24 CHOp24, γ-p24 Logjam and δ-p24 Baiser, all driven by *Cg-GAL4*. Fat body was dissected from wild type larvae and larvae in which genes encoding Ergic53, KdelR, Grasp65 and Tango1 had been knocked down under control of *Cg-GAL4*. Nuclei stained with DAPI (blue). Magnified insets in the lower right corner of each image show isolated GFP signal in white. Distribution of p24 proteins illustrated in bottom cartoons. See also Figure S2.

Next, we used co-immunoprecipitation followed by Western blotting to investigate the possibility that p24 proteins interacted with Tango1, required for their localization. We were in this way able to detect Tango1 when we immunoprecipitated GFP-tagged versions of Logjam (γ) and, to a lesser extent, of Eclair (α), CHOp24 (β) and Baiser (δ) from fat body adipocytes (Fig. 6 A). Because of the short length of the cytoplasmic tails of the p24 proteins (10 to 14 amino acid residues), we hypothesized that an interaction with Tango1 would most likely involve the ER luminal part of the protein, where the conserved GOLD domain is found. To test this, we expressed in the fat body a GFP-tagged version of Logjam from which we had deleted its GOLD domain (GFP-Logjam.ΔGOLD) and found that GOLD deletion abolished its interaction with Tango1 (Fig. 6 B). Similarly, deletion of the ER lumenal SH3 domain from a GFP-tagged version of Tango1 (Tango1.ΔSH3-GFP) prevented interaction with endogenous FLAG-tagged Logjam (Fig. 6 C). In addition to these co-immunoprecipitation experiments, we monitored the localization of GFP-Logjam.ΔGOLD and Tango1.ΔSH3-GFP, and found in both cases that the truncated proteins failed to localize correctly. In the case of Logjam, GOLD deletion resulted in ER and plasma membrane localization (Fig. 6 D), similar to the effect of Tango1 knock down (see Fig. 5). For Tango1, deleting its SH3 domain produced strong presence of the protein in the plasma membrane (Fig. 6 E), suggesting its scape from ERES. Altogether, these results show that Tango1 and p24 proteins interact through their respective SH3 and GOLD lumenal domains, which are required for the correct localization of both.

**Figure 6.**
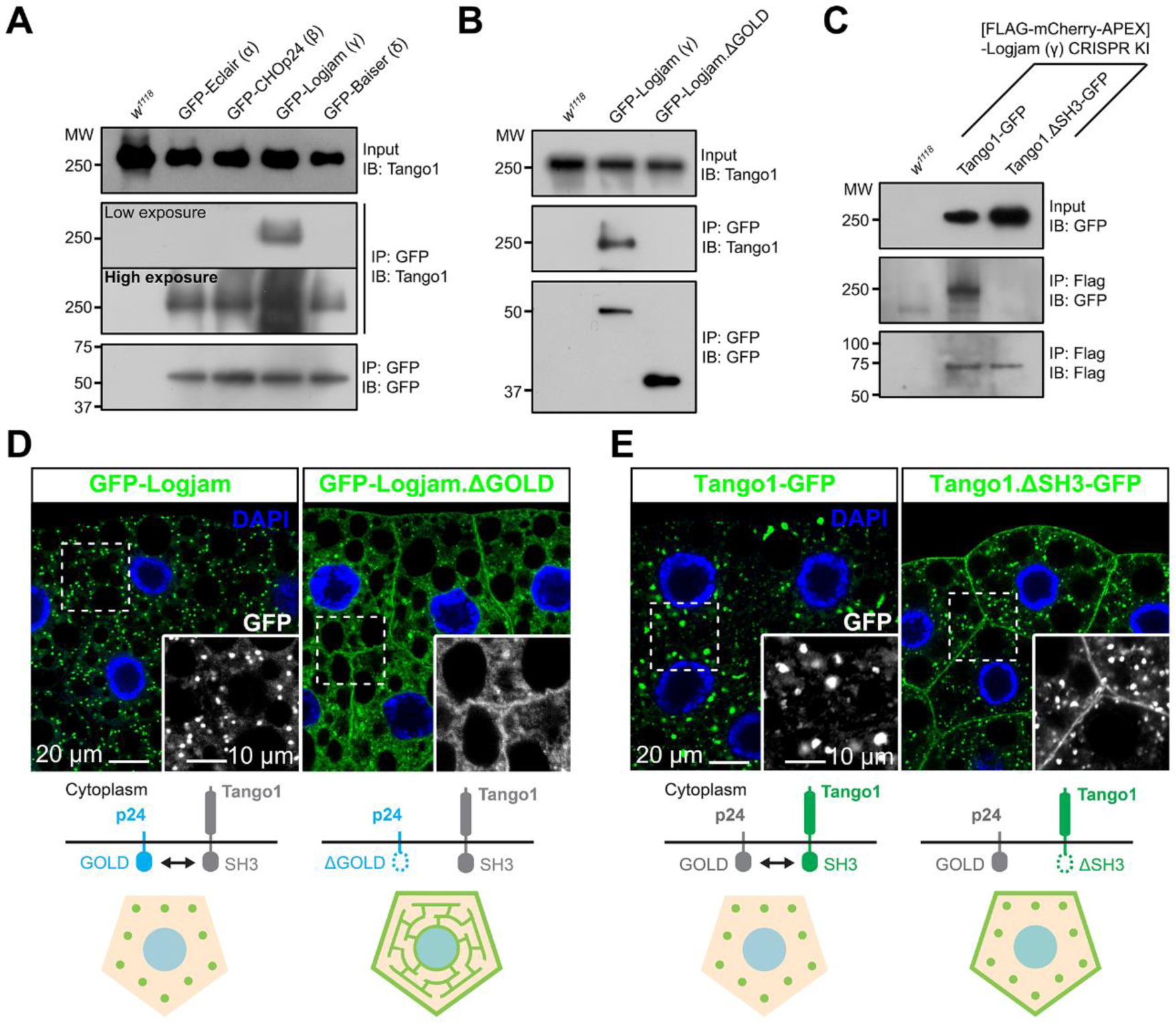
Tango1 and γ-p24 Logjam interact through their respective SH3 and GOLD domains. **(A)** Immunoblot analysis of Tango1-p24 interaction. GFP-tagged α-p24 Eclair, β-p24 CHOp24, γ-p24 Logjam and δ-p24 Baiser expressed under *Cg-GAL4* control were immunoprecipitated (IP) from L3 fat body lysates and immunoblotted (IB) with anti-Tango1 (both low and high exposure images are shown). **(B)** Immunoblot analysis of Tango1-Logjam interaction. Full-length and GOLD domain-deleted Logjam (Logjam.ΔGOLD), both GFP-tagged and expressed under *Cg-GAL4* control, were immunoprecipitated (IP) from L3 fat body lysates and immunoblotted (IB) with anti-Tango1. **(C)** Immunoblot analysis of Tango1-Logjam interaction. [FLAG]Logjam (CRISPR/Cas9 knock-in) was immunoprecipitated (IP) from L3 fat body lysates and immunoblotted (IB) with anti-GFP to detect full-length Tango1 and SH3 domain-deleted Tango1 (Tango1.ΔSH3), both GFP-tagged and expressed under *Cg-GAL4* control. As controls, *w^1118^* fat body was processed in parallel (A-C), and lysates and immunoprecipitates were immunoblotted, respectively, with anti-Tango1 and anti-GFP (A, B) or anti-GFP and anti-Flag (C). Uncropped scans are provided in Figure S5. **(D, E)** Confocal images of L3 fat body adipocytes showing in green localization of full-length and GOLD-deleted γ-p24 Logjam (D), and full-length and SH3-deleted Tango1 (E), all GFP-tagged and driven by *Cg-GAL4*. Magnified insets in the lower right corner of each image show isolated GFP signal in white. Nuclei stained with DAPI (blue). Protein interactions and distribution patterns of Logjam (D) and Tango1 (E) are schematically illustrated at the bottom.

### Maintenance of Tango1 at the ER-Golgi interface requires p24 proteins

Our experiments, revealing that Tango1 is required for localization of p24 proteins, additionally suggest that the converse is true as well, as hinted by Tango1.ΔSH3 mislocalization. To confirm the requirement of p24 proteins in Tango1 localization, we examined the localization of a GFP-tagged version of Tango1 upon knock down of p24 proteins. We found that knock down of *eclair* (α), *CHOp24+CG3908* (β) or *baiser* (δ) resulted in mislocalization of Tango1 to the plasma membrane (Fig. 7 A). Mislocalization of endogenous Tango1 could be detected as well with an antibody (Fig. S2 B). Interestingly, however, knock down of *logjam* (γ) failed to produce this effect (Fig. 7 A), suggesting that other p24 proteins, with which Tango1 interacts as well (Fig. 6 A), could compensate for the loss of Logjam to retain Tango1 at ERES. Consistent with this, co-immunoprecipitation experiments showed increased interaction between Tango1 and Eclair (α) (Fig. 7 B), CHOp24 (β) (Fig. 7 C) and Baiser (δ) (Fig. 7 D) when *logjam* expression was knocked down. From these results we conclude that p24 proteins prevent Tango1 escape from ERES. In addition, our data indicate that in their lumenal interaction with other proteins like Tango1, p24 subfamilies may show some functional redundancy (Fig. 7, E and F), in contrast to their non-redundant, interdependent requirements for localization.

**Figure 7.**
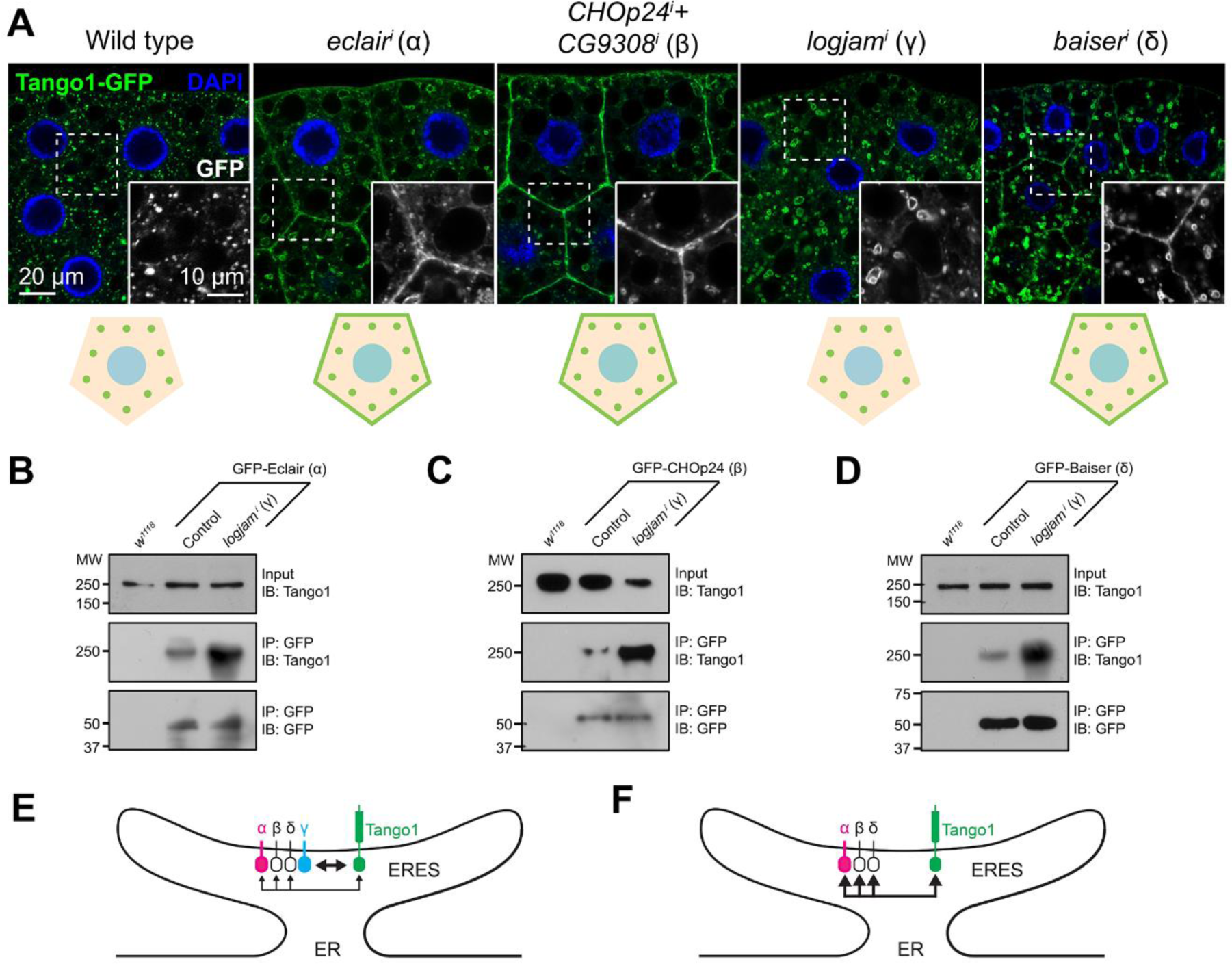
Loss of α-, β- or δ- but not γ-p24 causes Tango1 escape to the plasma membrane. **(A)** Confocal images of L3 fat body adipocytes showing in green localization of GFP-tagged Tango1 driven by *Cg-GAL4*. Fat body was dissected from wild type larvae and larvae in which genes encoding α-p24 Eclair, β-p24 CHOp24+CG9308, γ-p24 Logjam or δ-p24 Baiser had been knocked down. Magnified insets in the lower right corner of each image show isolated Tango1-GFP signal in white. Tango1 distribution patterns are schematically illustrated at the bottom. Nuclei stained with DAPI (blue). **(B-D)** Immunoblot analysis of Tango1-p24 interaction. GFP-tagged α-p24 Eclair (B), β-p24 CHOp24 (C) and δ-p24 Baiser (D), all expressed under *Cg-GAL4* control, were immunoprecipitated (IP) from control and *logjam^i^* L3 fat body lysates, and immunoblotted (IB) with anti-Tango1. As additional controls, *w^1118^* fat body was processed in parallel, and lysates and immunoprecipitates were immunoblotted with anti-Tango1 and anti-GFP, respectively. Uncropped scans are provided in Figure S5. **(E, F)** Schematic illustrations of Tango1-p24 interaction in presence (E) or absence (F) of γ-p24 Logjam. Arrow thickness represents interaction strength. See also Figure S2.

### Loss of p24 proteins expands COPII zone at ERES

To better understand the role of p24 proteins, and given their colocalization with COPII (Fig. 3, Q-T), we imaged endogenous GFP-tagged Sec13 (Sec13[GFP] knock-in) in the fat body upon knock down of *eclair* (α), *CHOp24*+*CG9308* (β), *logjam* (γ) or *baiser* (δ) (Fig. 8 A). In all four cases, Sec13 puncta in ERES-Golgi units exhibited a significant increase in their size and intensity (Fig. 8, B and C). Similarly, we could also detect an increase in the size and intensity of puncta formed by the COPII GTPase Sar1 (Fig. S3, A-C), and an enlargement of puncta positive for pre-*cis*-Golgi marker Grasp65 (Fig. S3, D-F), suggesting an expansion of this Golgi compartment. We have previously shown that in *Drosophila* ERES-Golgi units COPII concentrates in the center of ERES cups, whereas COPI displays a complementary localization around COPII, in the ERES periphery (Yang et al., 2021). To further characterize the alteration in COPII caused by the absence of p24 function, we used 3D-SIM to image simultaneously COPII coat component Sec13 and COPI coat component γCOP. When we knocked down *logjam* (γ) or *baiser* (δ), in contrast with the concentration of COPII contained to the center of wild type ERES, Sec13 signal expanded, partially overlapping peripheral COPI and adopting cup/doughnut morphologies typical of the latter (Fig. 8 D). Overall, these results demonstrate that p24 loss leads to an expansion of the COPII zone at ERES, strongly suggesting that p24 proteins serve an antagonistic role with respect to the COPII budding machinery.

**Figure 8.**
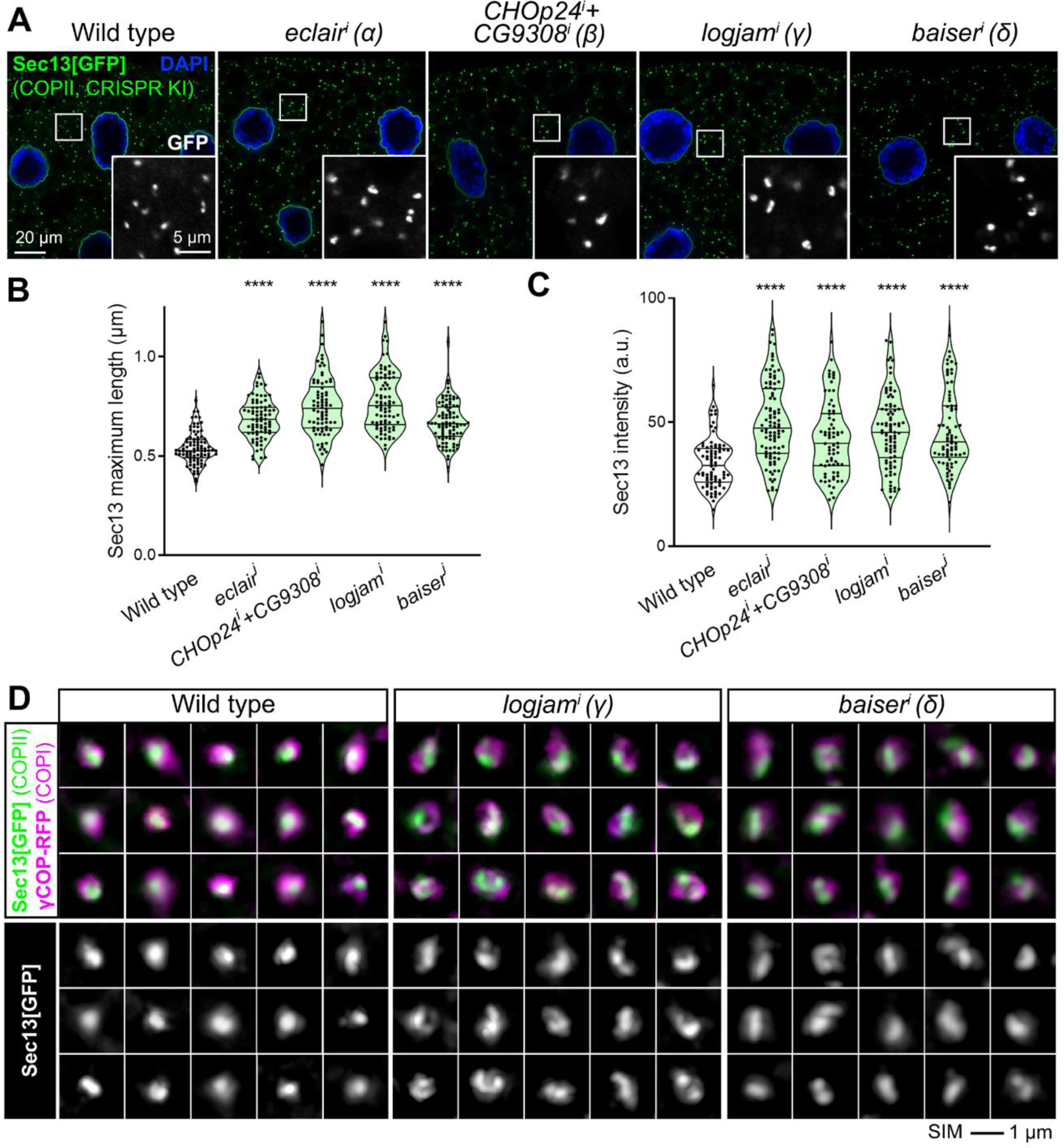
Loss of p24 increases COPII recruitment and expands COPII zone at ERES. **(A)** Confocal images of L3 fat body adipocytes showing in green localization of COPII coatomer Sec13[GFP] (CRISPR/Cas9 knock-in). Fat body was dissected from wild type larvae and larvae in which genes encoding α-p24 Eclair, β-p24 CHOp24+CG9308, γ-p24 Logjam or δ- p24 Baiser had been knocked down under control of *Cg-GAL4*. Magnified insets in the lower right corner of each image show isolated Sec13[GFP] signal in white. Tango1 distribution patterns are schematically illustrated at the bottom. Nuclei stained with DAPI (blue). **(B, C)** Quantification of maximum length (B) and intensity (C) of Sec13 puncta measured in images like those in A. Violin plots depict median value and interquartile range. Each dot represents a measurement in one punctum (n > 70 in each group). *p* values from Brown-Forsythe ANOVA and Dunnett’s multiple comparisons tests (B and C, *p* < 0.0001 (****) in all cases). **(D)** Super resolution 3D-SIM images of ERES-Golgi units from L3 fat body adipocytes showing localization of Sec13[GFP] (CRISPR/Cas9 knock-in, green) and γCOP-RFP (driven by *Cg-GAL4*, magenta). Fat body was dissected from wild type, *logjam^i^* and *baiser^i^* larvae (*Cg-GAL4*-driven knock down). Bottom images show isolated Sec13[GFP] signal in white. Images are maximum intensity projections of 3 to 5 sections. See also Figure S3.

### p24 proteins prevent excess vesicle budding

Intrigued by the observed expansion of COPII, we decided to further characterize the effect of p24 loss using FIB-SEM. To do this, we imaged with 20-nm z resolution volumes of wild type, *logjam*^i^ (γ) and *baiser^i^* (δ) fat body (two samples per genotype) and 3D-reconstructed ERES-Golgi units within them (ten units per genotype; Fig. S4 A). ERES, recognizable as regions of Golgi-facing ER devoid of ribosomes (Fig. 9 A), were reduced in size upon knock down of *baiser* (δ), but not *logjam* (γ) (Fig. 9, B and C). This is consistent with our earlier finding that Tango1 escapes ERES upon knock down of *baiser*, but not *logjam* (Fig. 7 A). Meanwhile, Golgi volume did not significantly change compared to the wild type (Fig. S4 B). We next analyzed tubular continuities we had previously discovered between ERES and pre-*cis*-Golgi (Yang et al., 2021), similar to ERES-ERGIC tubes others have independently described in cultured human cells (Shomron et al., 2021; Weigel et al., 2021), but found no difference in their frequency (around two per unit) across the three genotypes (Fig. S4, C and D). Besides tubular continuities, we identified between ERES and Golgi abundant vesicles in all three genotypes (Fig. 9 D). The number of these vesicles, however, showed a greater than two-fold increase in *logjam*^i^ (γ) and *baiser^i^* (δ) conditions compared with the wild type (Fig. 9 E). In the distribution of their sizes, vesicles from wild type, *logjam*^i^ (γ) and *baiser^i^* (δ) alike displayed a two-peaked diameter distribution, with peaks located at 52 nm and 64 nm (Fig. 9 F), consistent with COPI and COPII vesicle populations, respectively (Yang et al., 2021). When we separately analyzed vesicles by their diameter with a cut-off at 58 nm, the number of >58 nm vesicles increased in both *logjam*^i^ (γ) and *baiser^i^* (δ) ERES-Golgi units (Fig. 9 G). Furthermore, the added volume of >58 nm vesicles increased with respect to <58 nm vesicles in both *logjam*^i^ (γ) and *baiser^i^*(δ) (Fig. 9 H). We also analyzed the diameter of vesicular buds growing from ERES and Golgi (Fig. S4 E). In wild type ERES-Golgi units, same as in *logjam*^i^ (γ) and *baiser^i^* (δ), ERES buds were larger than Golgi buds, further supporting the existence of two populations of COPII and COPI vesicles at the ERES-Golgi interface in all three genotypes; at the same time, neither ERES nor Golgi buds significantly varied in diameter among the three genotypes (Fig. S4 F), indicating that p24 loss did not change their size. Finally, when we mapped the position of vesicles within the ERES cup, we observed that in *logjam*^i^ (γ) and *baiser^i^* (δ) more >58 nm vesicles were found in a now crowded peripheral zone (Fig. 9 H), reminiscent of our 3D-SIM data documenting COPII expansion (Fig. 8 D). In summary, our FIB-SEM analysis revealed a decrease in ERES size upon *baiser* (δ) knock down and an increase in the number of vesicles between ERES and pre-*cis*-Golgi after knock down of *baiser* (δ) or *logjam* (γ).

**Figure 9.**
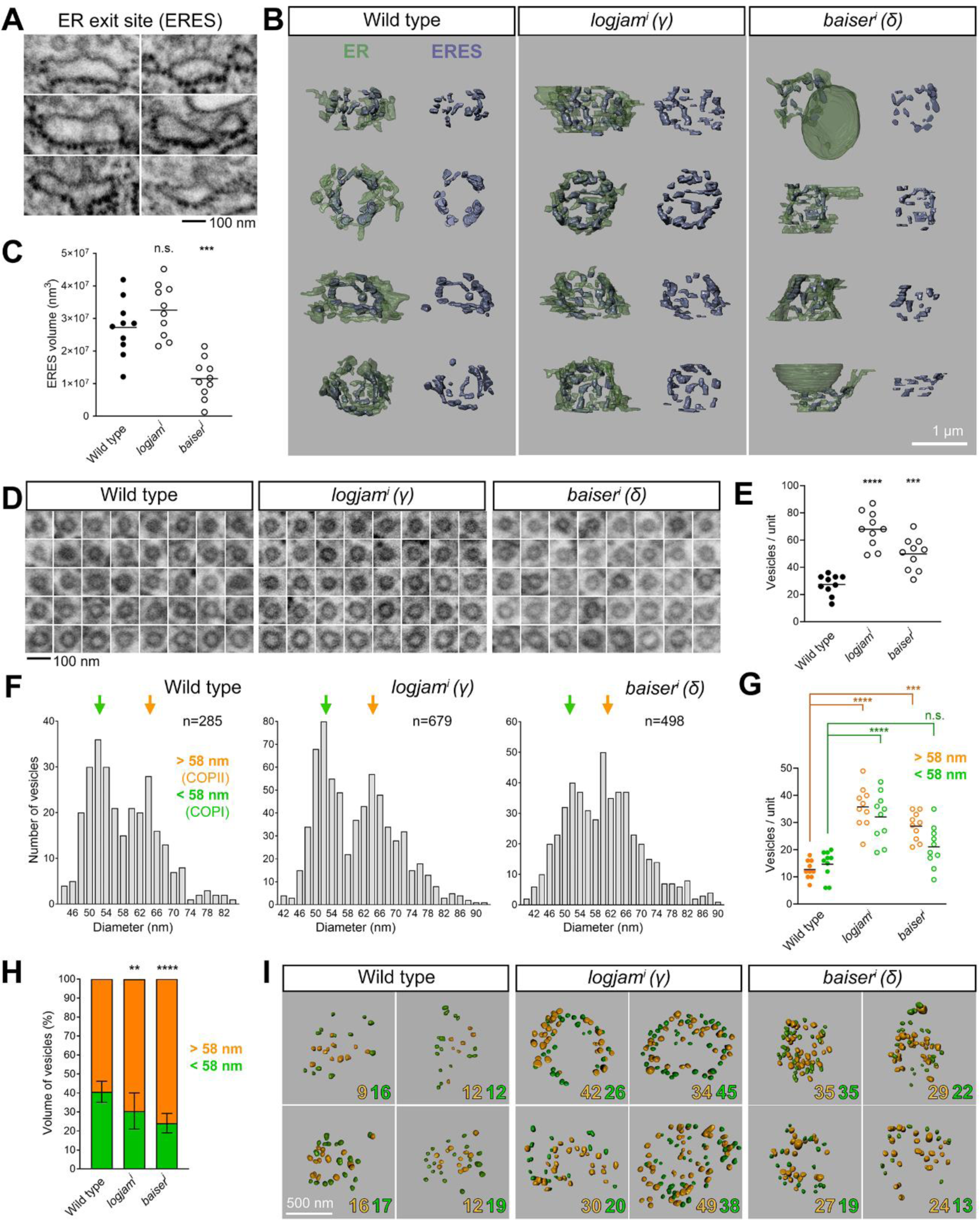
FIB-SEM analysis reveals ERES size reduction and increased vesicle budding upon p24 loss. **(A)** FIB-SEM images featuring examples of ERES areas, devoid of ribosomes on their Golgi-facing side. **(B)** 3D-reconstructions of ERES cups from FIB-SEM images of wild type, *logjam^i^* and *baiser^i^* L3 fat body adipocytes (knock-down driven by *BM-40-SPARC-GAL4*). Proper ERES (purple) shown separately from ER (green) on the right side. **(C)** ERES volume in wild type, *logjam^i^*and *baiser^i^* ERES-Golgi units. **(D)** FIB-SEM images exemplifying vesicles found between ERES and Golgi in wild type, *logjam^i^* and *baiser^i^* ERES-Golgi units. **(E)** Number of vesicles in wild type, *logjam^i^*and *baiser^i^* ERES-Golgi units. **(F)** Frequency distribution of apparent vesicle diameters in wild type, *logjam^i^* and *baiser^i^* ERES-Golgi units. Arrows indicate approximate peaks at 52 and 64 nm. **(G)** Number of vesicles larger (orange) and smaller (green) than a 58 nm diameter threshold in wild type, *logjam^i^* and *baiser^i^* ERES-Golgi units. Horizontal lines indicate mean value, with each dot representing one ERES-Golgi unit (C, E, G, n = 10 in each group). **(H)** Percentage of added vesicle volume corresponding to vesicles larger (orange) and smaller (green) than 58 nm in wild type, *logjam^i^* and *baiser^i^* ERES-Golgi units. Data represented as mean ± SD (n = 10 in each group). **(I)** Spatial distribution of vesicles larger (orange) and smaller (green) than 58 nm in wild type, *logjam^i^* and *baiser^i^* ERES-Golgi units. Counts for each annotated in the bottom right corner of 3D reconstructions. The plane of view in reconstructions is perpendicular to the *cis*-*trans* axis ERES-Golgi units (B and I). *p* values from one-way ANOVA and Dunnett’s multiple comparisons tests (C, *p* = 0.2282 (n.s.) for *logjam^i^* and 0.0002 (***) for *baiser^i^*; E, *p* < 0.0001 (****) for *logjam^i^*and = 0.0002 (***) for *baiser^i^*; G, *p* < 0.0001 (****) for *logjam^i^* >58 nm, = 0.0002 (***) for *baiser^i^*>58 nm, < 0.0001 for *logjam^i^* <58 nm and = 0.1102 (n.s.) for *baiser^i^* <58 nm; H, *p* = 0.0063 (**) for *logjam^i^*and < 0.0001 (****) for *baiser^i^*). See also Figure S4.

## DISCUSSION

In this study, we conducted a systematic characterization of p24 proteins in *Drosophila*, their role in the secretory pathway and the requirements for their localization. Our imaging of α-, β-, γ- and δ-p24 subfamily proteins showed that they concentrate between the ERES and pre-*cis*-Golgi, consistent with constant cycling between ER and Golgi. To maintain their localization, besides interactions among different p24 subfamilies (see below), ERES protein Tango1 is required. In absence of Tango1, p24 proteins fail to concentrate at the ER-Golgi interface and a fraction appears at the plasma membrane. Our further investigation of this relation revealed that Tango1 and p24 proteins physically interact and that this interaction involves the p24 GOLD domain and the SH3 domain of Tango1, both located in the ER lumen. Interestingly, the relation between p24 proteins and Tango1 is mutual, as Tango1 requires p24 presence as well to localize to ERES. Loss of p24 proteins α-, β-, or δ-p24 results in Tango1 mislocalization to the plasma membrane, whereas our results indicate that γ-p24 loss can be compensated by the other three. Overall, our results demonstrate that Tango1-p24 interplay is fundamental for maintaining a stable ER-Golgi interface (Fig. 10).

**Figure 10.**
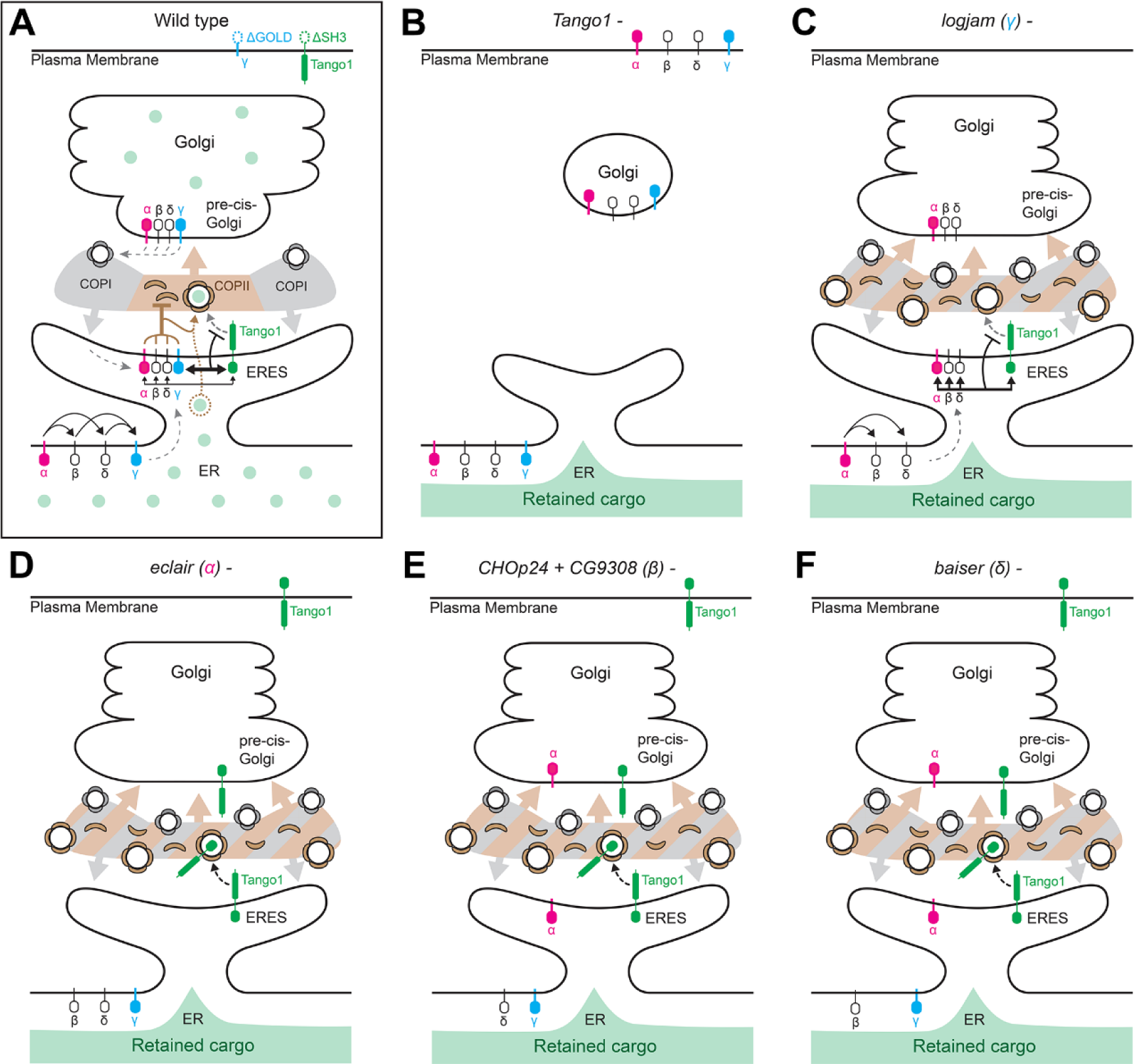
Tango1-p24 interplay at the ER-Golgi interface. **(A-F)** Schematic models depicting localization, interactions and roles of Tango1 and p24 proteins of the α-, β-, γ- and δ-p24 subfamilies in the wild type (A) and in conditions where Tango1 (B), γ- (C), α- (D), β- (E) or δ-p24 (F) proteins are absent. Concentration of p24 proteins from ER to ERES in the wild type (A) follows an α→βδ→γ hyerarchy of mutual requirements, possibly reflecting the assembly sequence of a heterotetramer. In this hierarchy, concentration of β- and δ- require presence first of α-p24 (D) and of each other (E, F), while γ-p24 requires all other three subfamilies. Once complexed at ERES, p24 proteins start cycling between ERES and pre-*cis*-Golgi transported by COPII (ER-to-Golgi) and COPI (Golgi-to-ER) vesicles (A). Interaction between the p24 GOLD domain (preferentially that of γ-p24) and the SH3 domain of Tango1 aids their concentration at the ER-Golgi interface (A). In the absence of Tango1 (B), uncoupling ERES from Golgi (Liu et al., 2017), p24 proteins are found in both ER and plasma membrane. Conversely, localization of Tango1 at ERES is dependent on p24 proteins, as in the absence of α- (D), β- (E) or δ-p24 (F), but not terminal γ-p24 (C), Tango1 leaves ERES and is trafficked forward to the plasma membrane. Apart from their effects on Tango1, p24 proteins of all four subfamilies are required for efficient general secretion, as in their absence all cargos we examined were retained in the ER (C-F). This is accompanied by an increase in COPII concentration, excess vesicle budding and expansion of the central COPII zone at ERES, all evidence of a negative role of p24 proteins on the COPII machinery. To reconcile secretory defects with increased COPII activity, we propose that p24 proteins act as concentrating receptors and ERES stabilizers, binding a wide range of cargos and other proteins like Tango1 to help their concentration at ERES while retarding their traffic forward.

To maintain the localization of p24 proteins, in addition, we were able to determine mutual requirements among the different p24 protein subfamilies. These requirements follow an α→β/δ→γ hierarchy in which α-p24 is capable of localizing to the ER-Golgi interface independently, β- and δ-p24 depend on the presence of α-p24 and of each other, and, lastly, γ-p24 needs the presence of all others. What determines these differential behaviors needs further investigation, since p24 proteins of all four subfamilies are very similar in sequence and organization. Due also to this similarity, it has been unclear whether p24 proteins play distinct roles or have overlapping functions. In this regard, our study strongly supports that p24 proteins of different subfamilies function non-redundantly, most probably as part of heterotetrameric complexes formed in the ER. Except for α-p24, localizing correctly by itself, complexing would be required for β-, γ- or δ-p24 entry into ERES or for initial concentration there, since they are found throughout the ER when mislocalized. In turn, γ-p24, for which the loss does not affect localization of any of the others, would be the last one to be incorporated into the putative heterotetramer. Consistent as well with the localization hierarchy we deduced, our phenotypic analysis demonstrates that p24 proteins are not functionally redundant across subfamilies, as members of all four subfamilies are required for efficient secretion. Our results, nonetheless, also argue that a partial complex lacking terminal γ-p24 retains some functionality. Indeed, loss of γ-p24 (Logjam) displays no Tango1 mislocalization. This is despite the fact that Logjam presents the strongest interaction with Tango1, showing that α-, β- and δ-p24 can cover for its loss. Therefore, while p24 proteins of the four subfamilies play non-redundant roles in the localization hierarchy, they may be able to bind redundantly through their GOLD domains Tango1 (and perhaps other proteins), albeit with different affinities.

From our findings here, a scenario emerges in which multiple mechanisms act on p24 proteins and Tango1 to maintain their localizations at the ER-Golgi interface. We propose that the balance of these forces results in a dynamic equilibrium that maintains a stable interface, ensuring its correct organization and efficient protein transport through it. For p24 proteins, forces influencing their localization are: (1) GOLD-SH3 interaction with Tango1, as in the absence of Tango1 all p24 proteins fail to concentrate at ERES and are instead found in both ER and plasma membrane; (2) interactions with p24 proteins of other subfamilies (except for α-p24) facilitates ERES concentration and may be required for their forward ER-Golgi transport, since p24 proteins mislocalized due to absence of others (β, γ and δ in absence of α; δ and γ in absence of β; β and γ in absence of δ) do not appear on the plasma membrane, and (3) retrograde transport that recycles p24 proteins from Golgi back to ERES, consistent with their well-attested binding to COPI (Dominguez et al., 1998; Fiedler et al., 1996). Meanwhile, Tango1 concentration at ERES depends on: (a) Tango1-Tango1 self-interaction through its cytoplasmic domains, consistent with our finding that the cytoplasmic part of the protein is sufficient to localize Tango1 to ERES (Liu et al., 2017); (b) SH3-GOLD interaction with p24 proteins, which impedes exit of Tango1 from ERES. In addition to these mechanisms, it has been reported that KDEL receptor-dependent Golgi-to-ERES recycling acts on both p24 proteins (Majoul et al., 2001) and Tango1 (Yuan et al., 2018). When we knocked down *Drosophila* KdelR, however, we did not see an effect on their localization. This may be due to qualitative differences across organisms or, alternatively, reflect that the influence of KdelR-dependent recycling of these proteins in *Drosophila* is not significant, its loss buffered by the added influence of other localization inputs. Nonetheless, our SIM imaging and previous visualization of Tango1 through APEX-TEM (Liu et al., 2017; Yang et al., 2021) seems to confirm that indeed concentration of *Drosophila* Tango1 at ERES does not result from its constant recycling from the Golgi but from a lack of forward transport from ERES.

Our functional characterization of p24 proteins, in addition, importantly uncovered a broad role for them in secretion. p24 proteins are widely regarded as specific transport receptors for ER-Golgi traffic of particular cargos such as GPI-anchored proteins (Muñiz et al., 2000; Schimmoller et al., 1995), WNT ligands (Port et al., 2011; Zang et al., 2015), insulin (Zhang and Volchuk, 2010) or Interleukin-1 (Zhang et al., 2020). In stark contrast, we show that knock down of p24 proteins in fat body adipocytes causes intracellular retention of all cargos we examined. Along with general secretion defects, we observed increased presence of COPII coatomer Sec13 and GTPase Sar1 at ERES, expanding their central COPII zone. At the same, FIB-SEM analysis revealed an excessive number of ERES-Golgi vesicles compared to the wild type. These defects were also seen after γ-p24 knock down, a condition in which Tango1 localization at ERES appears undisturbed. Therefore, general secretion impairments are likely caused directly by the loss of p24 proteins, rather than through Tango1 escape. Although the amount of COPII vesicles increases, the fact that general secretion is defective indicates that those vesicles do not mediate efficient transport. Our data, therefore, strongly support that p24 proteins negatively regulate the COPII machinery to prevent unproductive vesicle budding. Negative regulation of COPII vesicle biogenesis by p24 proteins agrees well with the fact that p24 mutations suppress Sec13 mutants in yeast (D’Arcangelo et al., 2015; Elrod-Erickson and Kaiser, 1996; Marzioch et al., 1999).

Expansion of the COPII zone and excess budding could be interpreted as a lax ER retention phenotype due to increased non-selective bulk flow (Gomez-Navarro et al., 2020; Lopez et al., 2020; Ma et al., 2017). p24 proteins, for instance, could take up space inside COPII vesicles, competing as decoys with non-cargo proteins for loading. However, arguing against this interpretation, cargos retained in fat body adipocytes upon p24 loss included secreted-GFP (GFP coupled to a secretion signal peptide), which should exit the ER through bulk secretion. Furthermore, for increased bulk secretion to result in more vesicles, as observed, bulk cargos should be able to recruit COPII and promote their own transport. An alternative explanation that would better fit our data is that p24 proteins function as concentrating receptors: through their lumenal GOLD domains, p24 proteins perhaps bind a broad range of cargos and are required to concentrate them at ERES, same as they are required for concentration of Tango1; meanwhile, through their cytoplasmic tails, p24 proteins could interact negatively with COPII to delay budding events and allow cargo loading. Interestingly, both Tango1 and p24 proteins bind COPII, and it has been proposed that Tango1 delays vesicle budding or excision to aid transport of cargos bound to its SH3 domain (Raote et al., 2018; Saito et al., 2009). Therefore, Tango1 and p24 proteins could have cooperative retardatory effects. In summary, we hypothesize that Tango1 and p24 proteins may both function as concentration receptors and ERES stabilizers (rather than as transport receptors or transport decoys) by binding an ample spectrum of cargos and other proteins in the ER lumen through their SH3 and GOLD domains, helping their concentration while retarding their exit from ERES.

Our study, finally, adds support to a central role of Tango1 in defining and maintaining ERES. Here, we proved a requirement of Tango1 in maintaining localization of p24 proteins through an SH3-GOLD domain interaction. From previous studies, Tango1 is known to interact through its cytoplasmic part with COPII (Saito et al., 2009), Syntaxin 18 (Nogueira et al., 2014), Rab1, Grasp65 (Liu et al., 2017), Sec16 and Sec12 (Maeda et al., 2017). Numerous proteins of the ER-Golgi interface, therefore, are coordinately bound by Tango1. Moreover, ERES reassembly after mitosis has been shown to depend on Tango1 (Maeda et al., 2020). Further supporting a structural role for Tango1 in the maintenance of the ER-Golgi interface, the loss of *Drosophila* Tango1 reduced the size of ERES and uncoupled them from Golgi, while overexpression of Tango1 created larger ERES (Liu et al., 2017; Yang et al., 2021). In light of all this evidence, we propose that Tango1 ensures building of a more stable ER-Golgi interface in animal cells through its multiple interactions, including lumenal binding to p24 proteins.

## Abbreviations

ERES: ER exit site
ERGIC: ER-Golgi intermediate compartment
FIB-SEM: Focused ion beam-scanning electron microscopy
GalT: Galactosyltransferase
GMAP: Golgi microtubule-associated protein
GOLD: Golgi dynamics
Grasp65: Golgi reassembly stacking protein of 65 kD
KdelR: KDEL receptor
ManII: Mannosidase II
SIM: Structured illumination microscopy
Tango1: Transport and Golgi organization 1

## ACKNOWLEDGMENTS

We thank Sally Horne-Badovinac for anti-Tango1 antibody and the Bloomington *Drosophila* Stock Center, Kyoto *Drosophila* Stock Center, Vienna *Drosophila* Resource Center and Tsinghua Fly Center for fly strains. We also thank the Tsinghua Center for Protein Research and Technology platform (Dr. Ying Li and Dr. Xiaomin Li) for technical help. This work was funded by grants 32150710524 and 91854207 from the National Natural Science Foundation of China and grant PID2021-122119NB-I00 from Ministerio de Ciencia e Innovación, all to J.C.P.-P. Research by J.C.P-.P. was also funded by the “Severo Ochoa” Program for Centers of Excellence (CEX2021-001165-S).

## AUTHOR CONTRIBUTIONS

K.Y. and F.Z conducted experiments. K.Y. and J.C.P.-P. analyzed the data and wrote the manuscript.

## MATERIALS AND METHODS

### Drosophila husbandry

Standard fly husbandry techniques and genetic methodologies, including balancers and dominant markers, were used to assess segregation of transgenes in the progeny of crosses, construct intermediate lines and obtain flies of the required genotypes for each experiment. Detailed genotypes in each experiment are provided in Table S1. Flies were cultured at 25°C in all experiments. The GAL4-UAS binary expression system was used to drive expression of UAS transgenes under temporal and spatial control of fat body GAL4 driver *Cg-GAL4* (2^nd^ chromosome) or *BM-40-SPARC-GAL4* (3^rd^ chromosome). Stable insertion of transgenic UAS constructs was achieved through standard P-element transposon transgenesis at Tsinghua Fly Center. Endogenous tagging was achieved through CRISPR/Cas9-assisted insertion (Peng et al., 2019) at Tsinghua Fly Center. The following strains were used:

*w^1118^* (BDSC:3605)

*w ; Cg-GAL4* (BDSC:7011)

*w ; BM-40-SPARC-GAL4 UAS-Dcr2/TM6B* (Liu et al., 2017)

*w ; UAS-eclair.RNAi^101388/KK^* (VDRC:101388)

*y sc v ; UAS-logjam.RNAi^HMS06058^* (THFC:TH04039.N)

*y sc v ; UAS-CHOp24.RNAi^HMC05582^* (THFC:TH04235.N)

*y sc v ; UAS-p24-1.RNAi^HMC04970^*(THFC:TH04238.N)

*y sc v ; UAS-p24-2.RNAi^HMS02005^* (THFC:THU4082)

*w ; UAS-CG9308.RNAi^6606/GD^* (VDRC:6606)

*w ; UAS-CG31787.RNAi^6372/GD^* (VDRC: 6372)

*y sc v sev ; UAS-opossum.RNAi^HMC02679^* (BDSC:43280)

*w ; UAS-baiser.RNAi^100612/KK^* (VDRC:100612)

*y w ; vkg^G454^-GFP/CyO* (DGRC:11069)

*w ; UAS-myr-RFP* (BDSC:7118)

*y w ; Rfabg-sGFP^fTRG.900^*(VDRC:318255)

*w ; UAS-GFP-GPI/(CyO)* (Greco et al., 2001)

*w ; UAS-mCD8-GFP/CyO* (BDSC:5137)

*w ; UAS-secr-GFP* (Pfeiffer et al., 2000)

*w ; UAS-GFP-KDEL* (BDSC:9898)

*y sc v sev ; UAS-KdelR.RNAi^HMC05779^* (BDSC:64906)

*w ; UAS-ManII.TagRFP* (BDSC:65249)

*w ; UAS-GFP-Eclair* (This study)

*w ; UAS-GFP-CHOp24* (This study)

*w ; UAS-GFP-Logjam* (This study)

*w ; UAS-GFP-Baiser* (This study)

*w ; [mCherry-APEX-Flag]Logjam* (This study)

*w ; UAS-GalT-TagRFP; TM2 / TM6B,Tb* (BDSC:65251)

*w ; UAS-ManII-EGFP; TM2 / TM6B,Tb* (BDSC:65248)

*w Gmap^KM102^-GFP* (DGRC:109702)

*w ; Grasp65[GFP-APEX-FLAG]* (This study)

*w ; Sec13[GFP-APEX-FLAG]* (This study)

*y sc v sev ; UAS-Grasp65.RNAi^HMC05584^* (BDSC:64565)

*w ; UAS-Ergic53.RNAi^108445/KK^*(VDRC:108445)

*w ; UAS-Tango1.RNAi^NIG11098R^/TM6B* (NIG:11098R)

*w ; UAS-Tango1-GFP* (Liu et al., 2017)

*w ; UAS-Tango1.*Δ*SH3-GFP* (This study)

*w ; UAS-GFP-Logjam.*Δ*GOLD* (This study)

*w ; UAS-Sar1-GFP-APEX* (Yang et al., 2021)

*y w ; Kr^If-1^ / CyO ; UAS-γCOP-mRFP* (BDSC:29714)

### Transgenic constructs

#### UAS-GFP-Eclair, UAS-GFP-CHOp24, UAS-GFP-Logjam, UAS-GFP-Baiser and UAS-GFP-Logjam.ΔGOLD

To produce each construct, the coding sequence of each gene was amplified from whole larva cDNA using PrimeScript RT-PCR Kit (Takara, cat # RR014-A). The amplified sequence was then purified through gel extraction (Magen HiPure Gel Pure DNA Mini kit, cat # D2111-03). Flanking att sequences were added through another round of PCR (Takara, cat # R011) and purified. The resulting products were then recombined into pDONR221 (Thermo Fisher Scientific, cat # 12536017) through a Gateway BP reaction with Gateway BP Clonase II Enzyme Mix (Thermo Fisher Scientific, cat # 11789020) to produce pDONR221-p24 entry clones. From there, p24 sequences were transferred into modified Gateway destination vector pTSGW (UASt-Signal peptide of Tango1-GFP-Gateway cassette) (Yang et al., 2021) through Gateway LR recombination using LR Clonase II Plus enzyme (Thermo Fisher Scientific, cat # 12538120) to obtain the desired plasmids.

Primers used were: Eclair-F, Eclair-R, att-Eclair-F and att-Eclair-R; CHOp24-F, CHOp24-R, att-CHOp24-F and att-CHOp24-R; Logjam-F, Logjam-R, att-Logjam-F, and att-Logjam-R; Baiser-F, Baiser-R, att-Baiser-F, and att-Baiser-R; LogjamΔGOLD-F, LogjamΔGOLD-R, att-Logjam-F, and att-Logjam-R. Primer sequences are listed in Table S2.

#### UAS-Tango1.ΔSH3-GFP

pDONR-Tango1ΔSH3 was generated through deletion PCR from pDONR-Tango1 (Liu et al., 2017) with primers Tango1ΔSH3-F and Tango1ΔSH3-R, and from there transferred into pTWG (UASt-Gateway cassette-GFP, Drosophila Carnegie Vector Collection) through Gateway LR recombination using LR Clonase II Plus enzyme. Primer sequences are listed in Table S2.

### CRISPR knock-in of [mCherry-APEX-FLAG]Logjam, Grasp65[GFP-APEX-FLAG] and Sec13[GFP-APEX-FLAG]

For knock-in of each gene, three plasmids were used: pU57-Donor-(gene of interest), pU6b-sgRNA-(gene of interest) and universal pU6b-sgRNA1. pU57-Donor consists of universal sgRNA1 sequence, 200 bp upstream sequence from the target site, tagging sequence, linker, 200 bp downstream sequence from the target site and sgRNA1 sequence. The target site of Logjam was right after its signal peptide sequence, while for Grasp65 and Sec13 it was C-terminal before their stop codons. pU57-Donor-Logjam, pU57-Donor-Grasp65 and pU57-Donor-Sec13 were synthesized by TsingKe Biotechnology Co.,Ltd.

For preparing pU6b-sgRNA for each gene, sgRNAs were selected on the website http://targetfinder.flycrispr.neuro.brown.edu and oligos were synthesized with TTCG and AAAC added at 5’ end of forward and reverse chain respectively. Then, sgRNA oligos were annealed and phosphorylated with T4 PNK (NEW ENGLAND BioLabs, cat # M0201) and T4 ligase buffer (NEW ENGLAND BioLabs, cat # M0202V). Next, sgRNA oligos were cloned into pU6b through a BbSI restriction enzyme site (NEW ENGLAND BioLabs, cat # R0539V) and T4 DNA ligation (NEW ENGLAND BioLabs, cat # M0202V) to obtain pU6b-sgRNA-Logjam, pU6b-sgRNA-Grasp65 and pU6b-sgRNA-Sec13. pU6b-sgRNA1 was prepared in the same way with a pair of universal sgRNA1 oligos.

The mixture of pU57-Donor-(gene of interest), pU6b-sgRNA-(gene of interest), and pU6b-sgRNA1 was injected to *y sc v ; nos-Cas9* embryos at Tsinghua Fly Center. Selected transgenic flies were all homozygous viable. Knock-in sites were validated by genome DNA sequencing. The detailed sequence of each component in pU57-Donor and the sequence of each sgRNA used are listed in Table S2.

### Confocal and 3D-SIM superresolution imaging

L3 larvae were predissected in PBS by turning them inside out with fine tip forceps, fixed in PBS containing 4% PFA (paraformaldehyde, Sinopharm Chemical Reagent, cat # 80096692) for 15 min, washed in PBS for 3 × 10 min, dissected from the carcass and mounted on a glass slide with a drop of DAPI-Vectashield (Vector Laboratories, cat # H-1200). Confocal images were acquired with a ZEISS LSM780 microscope equipped with a 63× oil Plan-Apochromat objective (NA 1.4) and a 100× oil Plan-Apochromat objective (NA 1.4).

SIM image stacks (z-steps of 0.24 μm) were acquired with a Nikon A1 N-SIM STORM microscope equipped with a CFI Apo SR TIRF 100× oil (NA 1.49) objective and an Andor Technology EMCCD camera (iXON DU-897 X-9255). Laser lines at 488, 560 and 640 nm were used for excitation. SIM image reconstructions were performed with NIS-Elements software (Nikon). Images are maximum intensity projections of three to five sections.

### Immunohistochemistry

Antibody stainings was performed using standard procedures for larval tissues. Briefly, larvae were predissected in PBS, fixed in PBS containing 4% PFA for 15 min, washed in PBS for 3 × 10 min, blocked in PBT-BSA (PBS containing 0.1% Triton X-100 detergent (Sigma-Aldrich, cat # T8787), 1% BSA (Zhongkekeao, cat # 201903A28) and 250 mM NaCl), incubated overnight with primary antibody in PBT-BSA in 4°C on a rotator. Next day, tissues were washed in PBT-BSA for 3 × 20 min, incubated for 2 h with secondary antibody in PBT-BSA at room temperature, washed in PBT-BSA for 3 × 20 min and then PBS for 3 × 10 min. Fat body tissues were finally dissected and mounted on a glass slide with DAPI-Vectashield. The primary antibody guinea pig anti-Tango1 (Lerner et al., 2013) (1:1000) was used. Secondary antibodies were goat anti-guinea pig IgG (1: 200, Alexa Fluor 488 conjugated, Jackson ImmunoResearch, cat # 106545003; 1:200, Alexa Fluor 647 conjugated, Jackson ImmunoResearch, cat # 106605003).

### Immunoprecipitation

L3 fat body from 200 larvae was collected and homogenized on ice using an electric pellet pestle and a lysis buffer containing 10 mM Tris-HCl (pH=7.5), 0.5 mM EDTA, 150 mM NaCl, 0.5% NP-40 (Leagene, cat # 1221A21) and 1x protease inhibitor (Beyotime, cat # P1005). Samples were then cleared through centrifuging for 15 min at 20,000 g and 4 °C. Protein concentration of lysates was quantified using a BCA kit (Thermo-fisher, cat # 23227) with a NanoDrop 2000C. GFP and Flag immunoprecipitation experiments were then conducted according to manufacturer’s instructions.

For GFP immunoprecipitation, GFP-Trap agarose beads (ChromoTek, cat # GT10) were first washed with IP buffer (10 mM Tris-HCl, 150 mM NaCl, 0.5 mM EDTA, pH = 7.5) for 3 × 1 min and collected through centrifuging for 1 min at 5,000 g, 4 °C. Samples were incubated with pre-washed GFP agarose beads and rotated overnight at 4 °C. Then, beads were collected through centrifuging for 1 min at 5000 g, 4 °C. Next, beads were washed 5 × 1 min with IP buffer, and collected by centrifuging for 1 min at 5,000 g, 4 °C. Finally, 2 × SDS-PAGE buffer (120 mM Tris-HCl, 4% SDS, 20% glycerol, 0.5% bromophenol blue, and 5% β-mercaptoethanol) was added to samples, and proteins were eluted from GFP agarose beads through boiling at 95 °C for 10 min, and cooled down on ice.

For Flag immunoprecipitation, anti-Flag magnetic beads (Sigma-Aldrich, cat # M8823) were washed using 1 × TBS buffer (50 mM Tris-HCl, 150 mM NaCl, pH = 7.5) for 3 times and collected by a magnetic rack. Samples were incubated with pre-washed anti-Flag magnetic beads and rotated overnight at 4 °C. Then, beads were collected, washed for 5

× 1 min with 1 × TBS buffer. Proteins were eluted from Flag magnetic beads through adding 5 package of beads gel volume of 150 ng/μl 3 x Flag peptide (Sigma-Aldrich, cat # 4799) in 1 × TBS buffer and incubating in a rotator for 1 h at 4 °C. Then, supernatant was collected, added with 5 x SDS protein loading buffer (250 mM Tris-HCl, 10% SDS, 50% glycerol, 0.5% bromophenol blue and 5% β-mercaptoethanol), boiled at 95 °C for 10 min and cooled down on ice.

### Western blotting

Protein lysates added with 5 × SDS protein loading buffer were boiled at 95 °C for 10 min for reducing. Then, sample were loaded in a 4 - 20% SDS-PAGE gradient gel (Beyotime, cat # P0057A) or 15% SDS-PAGE gel (Beyotime, cat # P0055B) and separated by electrophoresis in 1 × SDS-PAGE running buffer (Beyotime, cat # P0014A) at 120 V. Proteins were then transferred to PVDF membrane (Bio-rad, cat # 1620177) for 70 min at 300 mA, and blocked in 5% skim milk in TBST (50 mM Tris-HCl, 150 mM NaCl, 0.5% Tween-20, pH = 8) for 1 h at room temperature. Primary antibodies were diluted in blocking solution and incubated overnight at 4 °C on a rotator. Next day, membranes were washed 3 × 10 min with TBST, incubated with secondary antibodies diluted in TBST for 1 h at room temperature on a rotator, washed 3 × 10 min with TBST and exposed with a ECL kit (Bio-rad, cat # 1705061) on X-ray films (Kodak, cat # JPKD-5). Primary antibodies used were: anti-Tango1 (Lerner et al., 2013) (1:5000), anti-GFP (1:3000, Roche, cat # 1814460001) and anti-Flag (1:3000, Sigma Aldrich, cat # F1084). Secondary antibodies used were: Goat anti-mouse IgG-HRP (1:10000, Abmart, cat # M21001L), Goat anti-Guinea pig IgG-HRP (1:10000, Abcam, cat # ab6908).

### FIB-SEM imaging

FIB-SEM imaging was performed as previously described (Yang et al., 2021). Resin blocks were trimmed to expose tissues and then fixed onto a 45/90° screw type holder. The samples were subsequently coated with a layer of gold using a HITACHI E-1010 ion sputter coater for 120 s. FIB-SEM imaging was performed using a FEI Helios NanoLab G3 dual beam microscope system equipped with ETD, TLC and ICD cameras (ThermoFisher Scientific). During the milling of slices, an ion beam current of 0.43 nA at a 30 kV acceleration voltage was employed, with each milling step set at 20 nm. For the SEM imaging, the following parameters were utilized: a beam current of 0.4 nA, an acceleration voltage of 2 kV, a working distance of 2 mm, a dwell time of 8 µs, a pixel size of 3 - 4 nm and a pixel count of 4096 × 3536. TLD and ICD cameras collected backscattered signal for imaging. Imaging software used was AutoSlice and View G3 1.7.2 (FEI).

Images obtained from FIB-SEM were imported into Dragonfly (Object Research Systems). The Dragonfly Image Loader was utilized to import the images and the SSD method in the Slide Registration panel was employed to align them. For segmentation, various Regions of Interest (ROIs) were created using the ROI tools panel. Each organelle or membrane component was manually segmented as an individual ROI using the ROI Painter round brush tool in 2D mode. Once segmentation was completed for each section, the ROIs were exported and saved as object files. These objects were then converted into 3D meshes by using the export box in the ROI Tools panel. The meshes underwent a smoothing process four to six times and were examined in 3D scene mode, appearing as solid and fully opaque objects. For volume measurements, the information panel was used to record values for each object. The diameter of each vesicle, visible in 2-4 consecutive sections, was measured on its largest xy section in 2D mode using the Ruler tool in the annotation panel.

### Statistical analysis

Fluorescence intensity, fluorescence profiles and puncta size were calculated using Image J. Statistical analysis and graphical representations were performed using GraphPad Prism. *p* values were calculated by one-way ANOVA and Dunnett’s multiple comparisons tests, Brown-Forsythe ANOVA and Dunnett’s multiple comparisons tests or unpaired t test. For all analysis, significance was determined at p < 0.05 (*p < 0.05, **p < 0.01, ***p < 0.001, ****p < 0.0001). Statistical details of each experiment are listed in the Figure legends.

---

**Figure S1.**
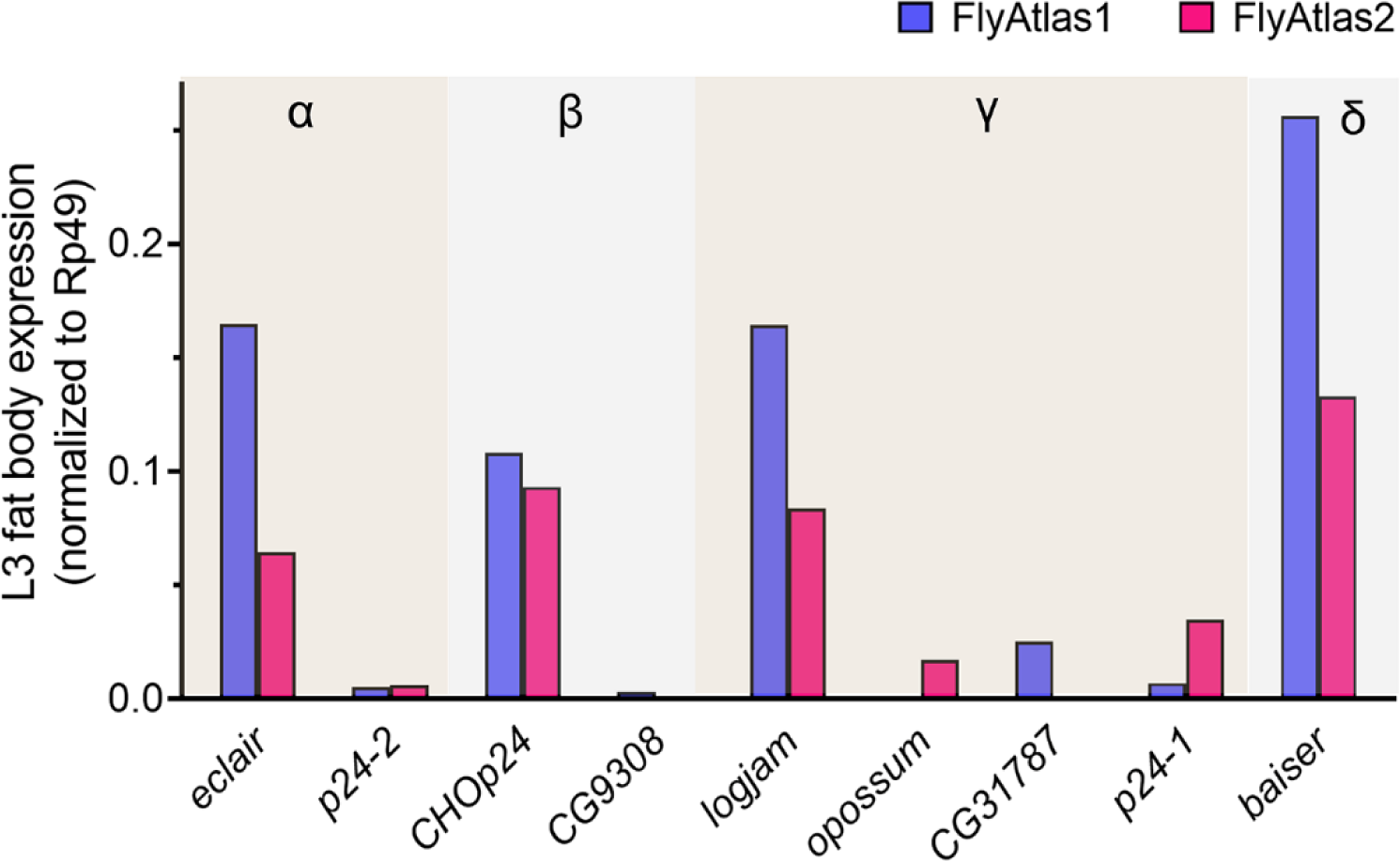
p24 expression level in larval fat body cells. Expression of genes encoding p24 family proteins in L3 fat body adipocytes according to microarray-based FlyAtlas1 data (http://flyatlas.org/atlas.cgi; violet) and RNAseq-based FlyAtlas2 data (https://flyatlas.gla.ac.uk/FlyAtlas2/; fuchsia). Expression levels are gene abundance normalized to that of *Rp49*. Related to Figure 1.

**Figure S2.**
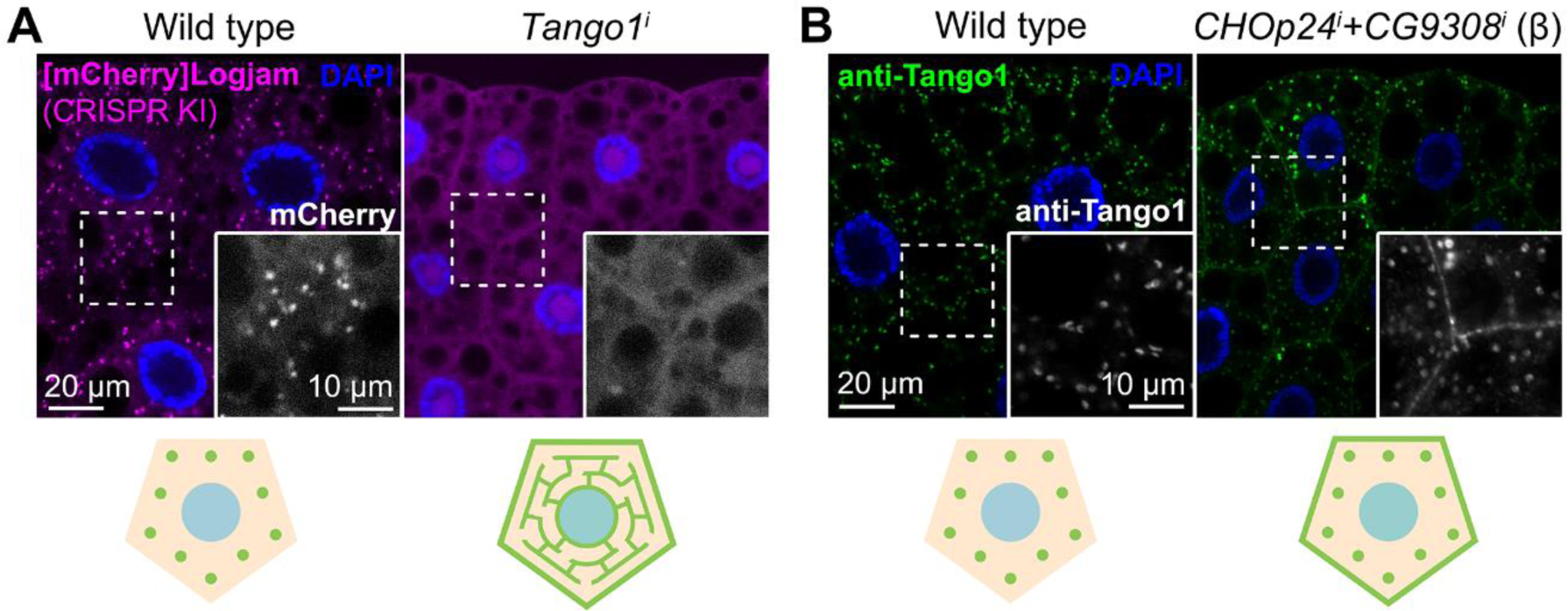
Tango1-p24 localizations are mutually dependent. **(A, B)** Confocal images of L3 fat body adipocytes showing localization of endogenous γ-p24 Logjam in magenta ([mCherry]Logjam CRISPR/Cas9 knock-in, A) and endogenous Tango1 in green (anti-Tango1, B). Fat body was dissected from wild type larvae (A, B), and larvae in which genes encoding Tango1 (A) or CHOp24 and CG9308 (B) had been knocked down under control of *Cg-GAL4* (A) or *BM-40-SPARC-GAL4* (B). Magnified insets in the lower right corner of each image show isolated Logjam (A) or Tango1 (B) signal in white. Nuclei stained with DAPI (blue). Observed distribution of proteins illustrated in bottom cartoons. Related to Figure 5 and Figure 7.

**Figure S3.**
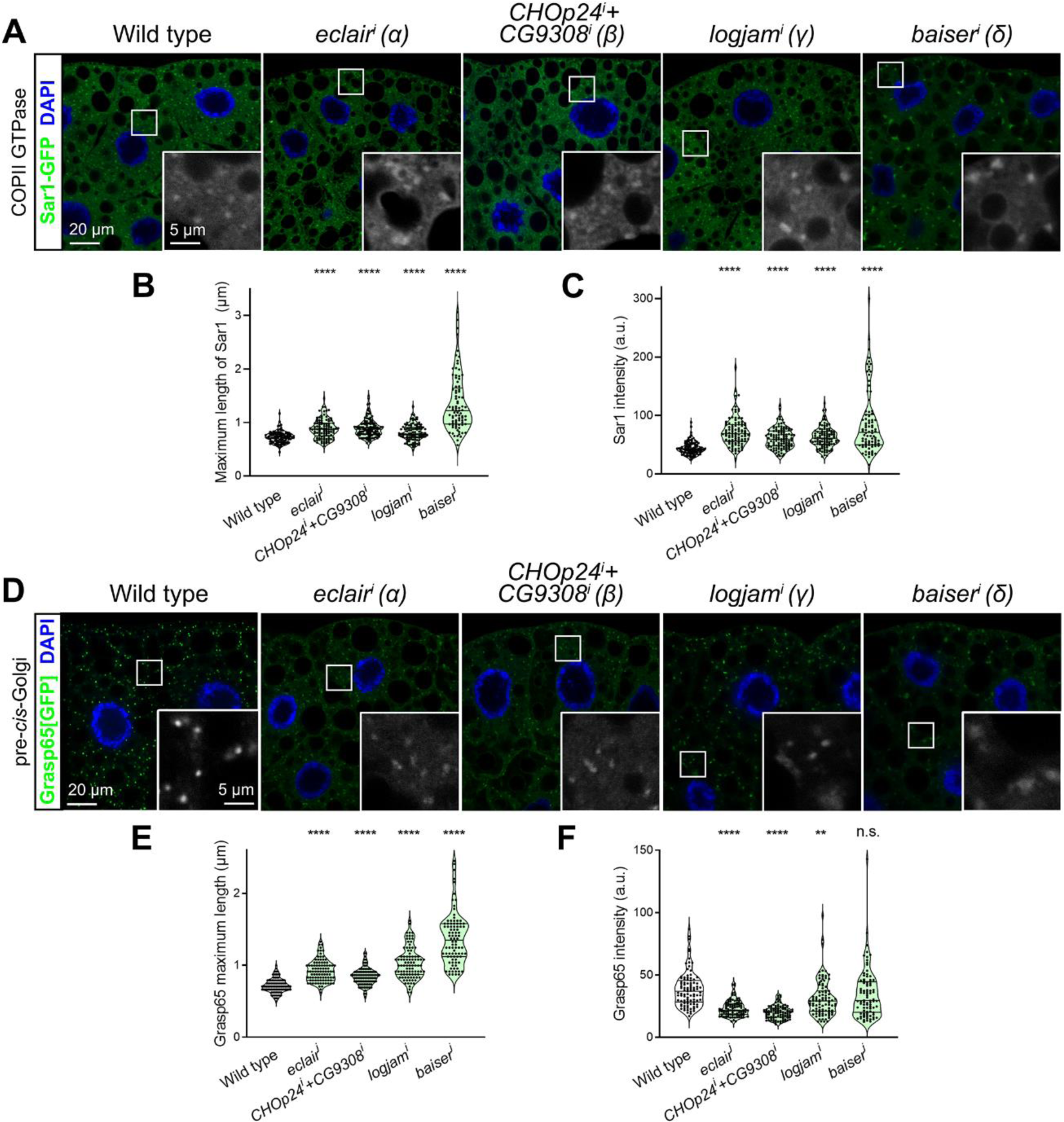
p24 loss increases Sar1 recruitment and enlarges pre-*cis*-Golgi. **(A, D)** Confocal images of L3 fat body adipocytes showing in green localization of COPII GTPase Sar1-GFP (driven by Cg-GAL4, A) and Grasp65[GFP] (CRISPR/Cas9 knock-in, D). Fat body was dissected from wild type larvae and larvae in which genes encoding α-p24 Eclair, β-p24 CHOp24+CG9308, γ-p24 Logjam or δ-p24 Baiser had been knocked down under control of *Cg-GAL4*. Magnified insets in the lower right corner of each image show isolated Sec13[GFP] signal in white. Tango1 distribution patterns are schematically illustrated at the bottom. Nuclei stained with DAPI (blue). **(B, C, E, F)** Quantification of maximum length (B, E) and intensity (C, F) of Sar1 puncta (B, C; measured in images like those in A) and Grasp65 puncta (E, F; measured in images like those in D). Violin plots depict median value and interquartile range. *p* values from Brown-Forsythe ANOVA and Dunnett’s multiple comparisons tests (B, C, E, *p* < 0.0001 (****) for *eclair^i^*, *CHOp24^i^+CG9308^i^*, *logjam^i^* and *baiser^i^*; F, *p* < 0.0001 (****) for *eclair^i^* and *CHOp24^i^+CG9308^i^*, = 0.0027 (**) for *logjam^i^*, and = 0.4924 (n.s.) for *baiser^i^*). Related to Figure 8.

**Figure S4.**
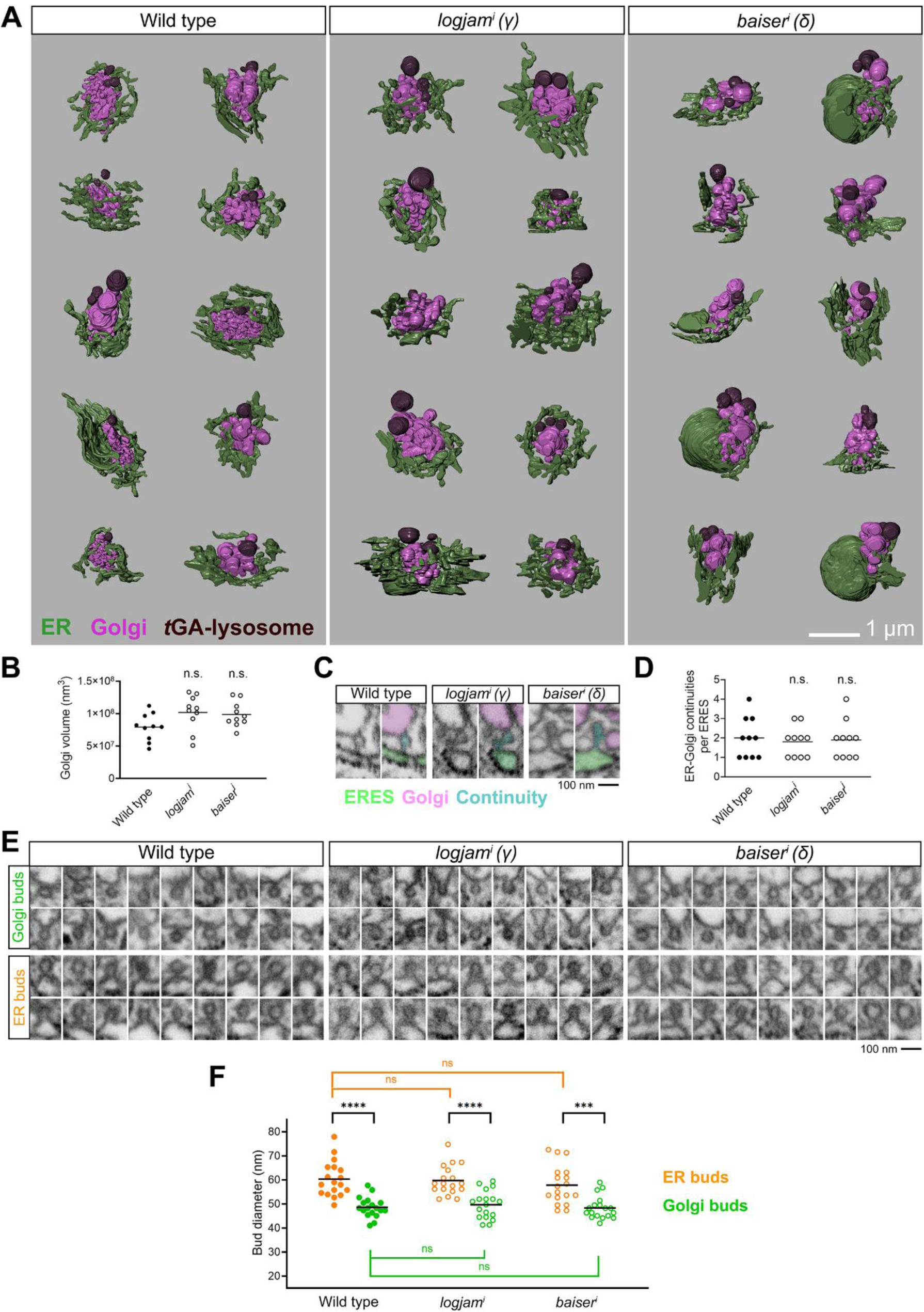
FIB-SEM analysis of mutant p24 ERES-Golgi units. **(A)** 3D-reconstructions of ERES-Golgi units from FIB-SEM images of wild type, *logjam^i^* and *baiser^i^* L3 fat body adipocytes (knock-down driven by *BM-40-SPARC-GAL4*). 10 ERES-Golgi units per genotype were reconstructed for our FIB-SEM analysis. Different colors indicate ER (green), Golgi (pink) and trans-Golgi associated (tGA) lysosomes (Zhou et al., 2023) (brown). **(B)** Golgi volume in wild type, *logjam^i^* and *baiser^i^*ERES-Golgi units. **(C)** FIB-SEM images of ERES-Golgi continuities in wild type, *logjam^i^* and *baiser^i^* ERES-Golgi units. Different colors in the color-coded version of each image indicate ERES (green), Golgi (pink) and continuity (cyan). **(D)** Number of ERES-Golgi continuities in wild type, *logjam^i^* and *baiser^i^* ERES-Golgi units. Each dot represents an ERES-Golgi unit (B and D, n = 10 in each group). **(E)** FIB-SEM images exemplifying vesicle buds found in ERES and pre-*cis*-Golgi in wild type, *logjam^i^* and *baiser^i^* ERES-Golgi units. **(F)** Quantification of the apparent diameter of ER (orange) and Golgi (green) buds in wild type, *logjam^i^* and *baiser^i^* ERES-Golgi units. Each dot represents one bud (n = 18 in each group). Horizontal lines indicate mean value (B, D, F). *p* values from one-way ANOVA and Dunnett’s multiple comparisons tests (B, *p* = 0.0622 (n.s.) for *logjam^i^* and 0.1162 (n.s.) for *baiser^i^*; D, *p* = 0.8636 (n.s.) for *logjam^i^* and 0.9605 (n.s.) for *baiser^i^*; F, *p* = 0.9655 (n.s.) for *logjam^i^*ER buds, 0.5630 (n.s.) for *baiser^i^* ER buds, 0.7925 (n.s.) for *logjam^i^* Golgi buds and 0.9905 (n.s.) for *baiser^i^*Golgi buds), Welch’s *t* test (F, *p* < 0.0001 (****) for ER v.s. Golgi buds in wild type, = 0.0002 (***) for ER v.s. Golgi buds in *baiser^i^*), and unpaired *t* test (F, *p* < 0.0001 (****) for ER v.s. Golgi buds in *logjam^i^*). Related to Figure 9.

**Figure S5.**
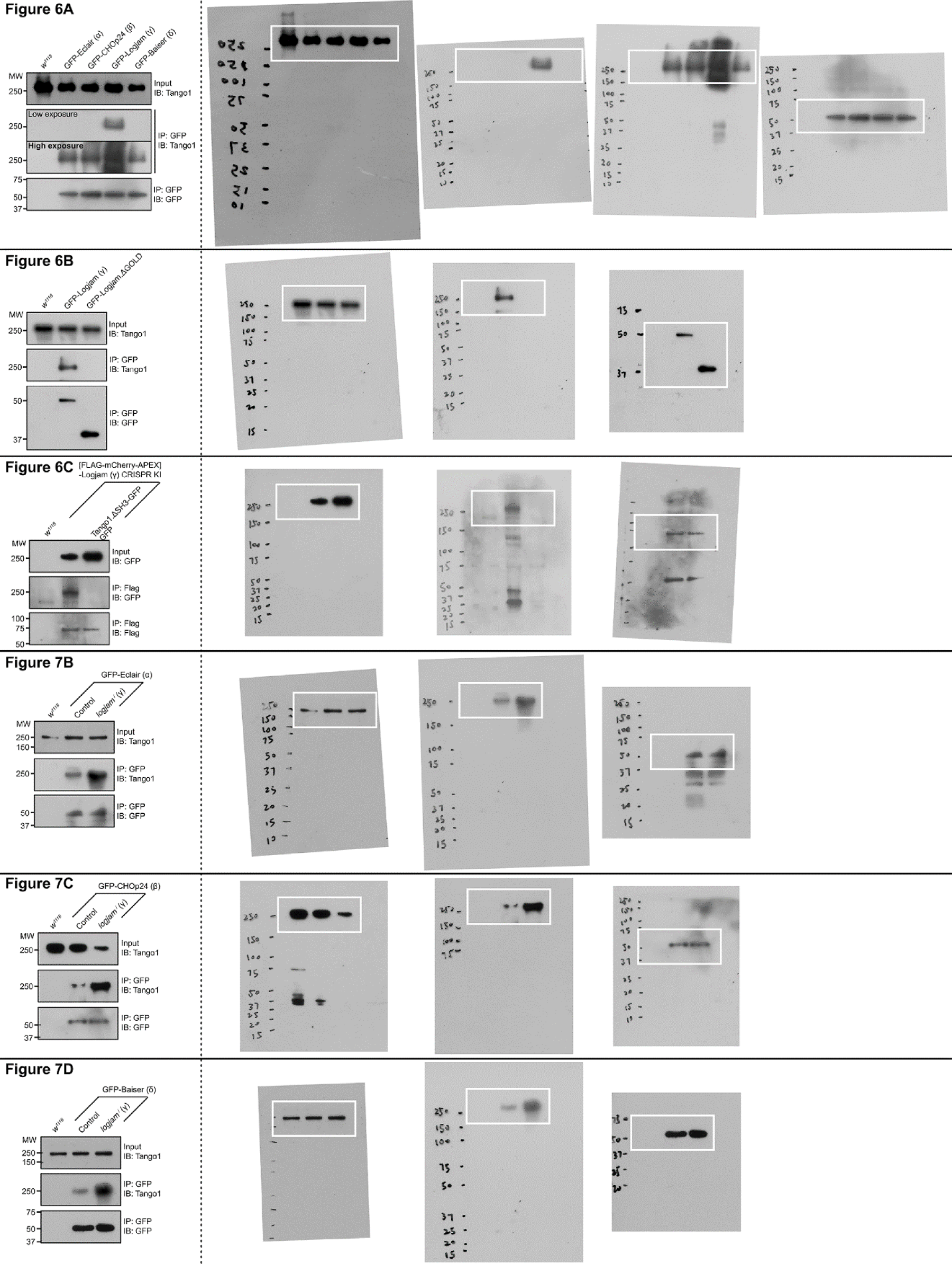
Uncropped Western blot scans.

